# A Stochastic Model of Metastatic Bottleneck Predicts Patient Outcome and Therapy Response

**DOI:** 10.1101/348086

**Authors:** Ewa Szczurek, Tyll Krueger, Barbara Klink, Niko Beerenwinkel

## Abstract

Metastases are responsible for 90% of cancer-related deaths. Initiation of metastases, where newly seeded tumor cells expand into colonies, presents a tremendous bottleneck to metastasis formation. Despite its clinical importance, our understanding of this process is very limited. Here, we propose a simple stochastic model assuming that the initiating metastatic cells proliferate faster when surrounded by more of their kind. The model quantifies the severity of metastatic bottleneck as the probability that the seeded colony survives. Based on this model, we derive how metastasis occurrence depends on primary tumor size and affects patient outcome. Our predictions agree with epidemiological data for thirteen cancer types. The model predicts that impact of treatment decisions depends both on the primary tumor size and on the severity of the metastatic bottleneck, and that medical interventions that tighten the bottleneck would be much more efficient than therapies that decrease overall tumor burden, such as chemotherapy.

## 1. Introduction

Metastasis formation is a multi-step process, in which tumor cells spread from the primary site and colonize distant organs (1). Metastases are responsible for 90% of deaths from cancer (2, 3), yet their formation is known to be highly inefficient (4–6). Metastasis formation and the bottleneck that this process encounters at initialization of metastatic colonies, is still poorly understood (7), both in terms of the underlying biological process and in terms of quantifying how severe the bottleneck is. Targeting the metastatic bottleneck would have a huge therapeutic potential for cancer, as it presents a natural point of attack of this fatal disease.

The process of metastasis formation can be divided into three phases (Figure 1a). The first phase consists of tumor cell entry into the vascular system (intravasation), transport in the blood, and exit from the vascular system to a secondary organ (extravasation). Presumably, this may be preceded by the acquisition of necessary genetic or epigenetic alterations in the primary tumor (6–8). Experimental data suggests that the tumor cells that are shed from the primary site are already equipped with metastatic abilities (9–11). Recent findings support that metastatic tumor cell dissemination begins in early (12, 13), rather than late (14) stage of the disease. The first phase is highly efficient, as the released tumor cells deal remarkably well with the obstacles of delivery to distant organs and their infiltration (6, 8, 15–17). In contrast, the second phase—metastasis initiation—is extremely inefficient (4–7). Relative to the huge numbers of cells that disseminate during the long period of primary tumor growth, only very few of them successfully form distant metastases (15, 18). This bottleneck is commonly understood as the lack of compatibility of the seeded tumor cells with the soil they encounter in the affected organ (6, 19, 20). In mice models, the metastatic seeding potential was observed to increase with the size of tumor cell clumps (21), which was recently confirmed for human circulating tumor cell clusters (22). Still, compared to how much is known about the dissemination and transport of tumor cells to the secondary organ, the mechanism behind the colonization bottleneck remains a mystery (7). In the last phase of metastasis formation, those successful colonies that do survive form micrometastases and, subsequently, macrometastases, which become clinically detectable (7).

**Figure 1.**
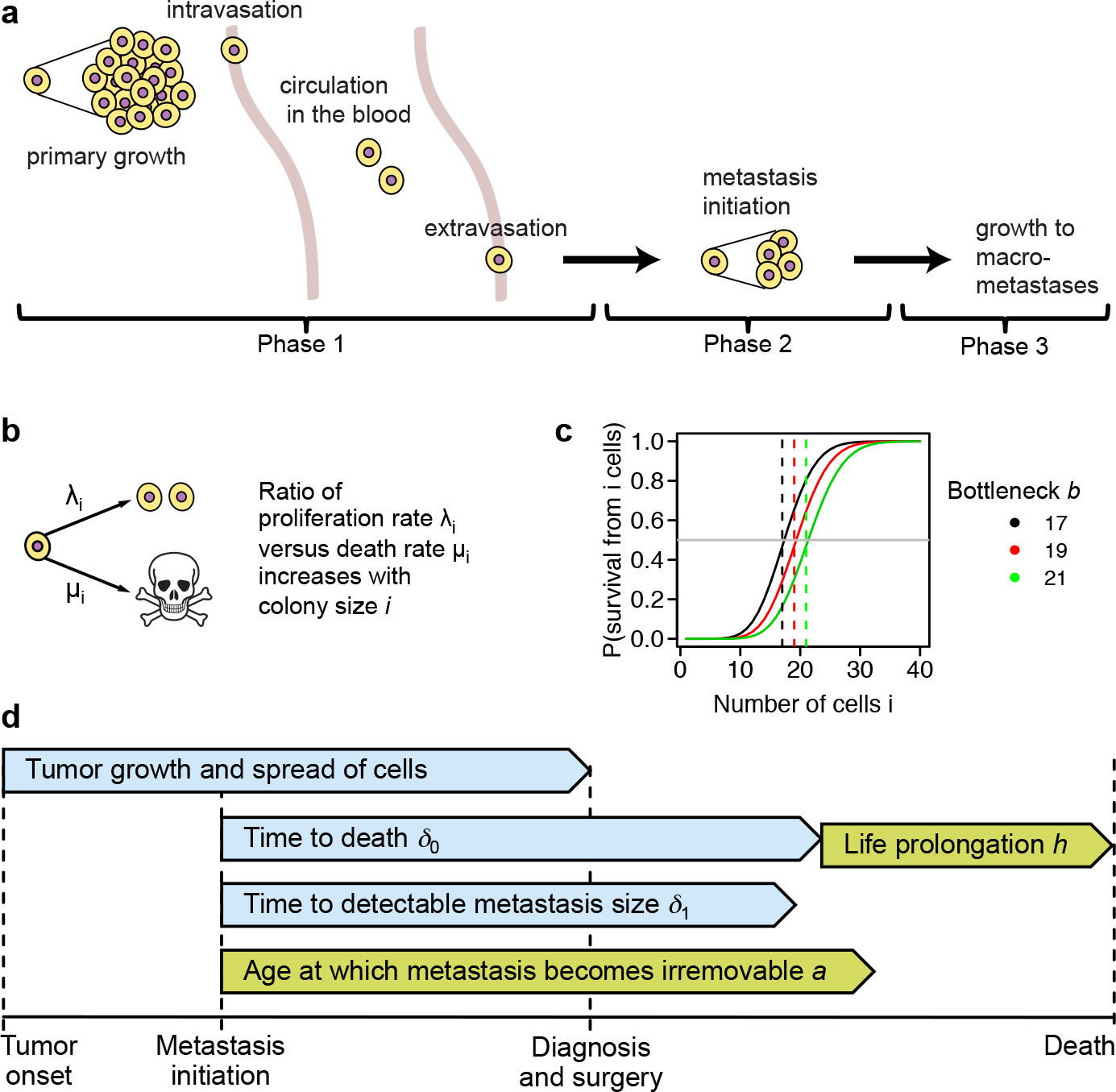
Modeling metastasis formation process and its bottleneck. **a** Three phases of metastasis formation, Phases 1 and 3 being very efficient. In Phase 1, tumor cells are constantly released into the blood stream. Given that larger tumors can release more cells over time, they have a higher chance to develop metastases. Thus, a dependency of metastasis probability and patient outcome on tumor size is expected. The bottleneck is the metastasis initiation in Phase 2. **b** Stochastic birth-and-death process of metastasis initiation. For each cell in the forming colony, the ratio of its proliferation rate λ_*i*_ to its death rate *μ*_*i*_ depends on the total number of cells *i* with a proportionality constant 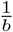. **c** The bottleneck severity *b* is the critical colony size for metastasis initiation. The probability of colony survival starting from *i* cells is lower than 0.5 (gray horizontal line) when *i* < *b* and larger than 0.5 when *i* > *b*, and increases rapidly when *i* crosses *b*. At the point when the colony has *b* cells, the jump probability that the colony will increase its size by one equals roughly 0.5. *b* = 19, red, is a median bottleneck commonly observed in our model; Figure 4) **d** Model parameters on a timescale. From tumor onset to diagnosis, the tumor grows and releases cells. Once a metastasis is successfully initiated, it takes a cancer-specific average number of years *δ*_0_ until it becomes lethal. The initiated metastases grow on average *δ*_1_ years to become large enough to be detectable by screening. With a certain probability, decreasing with metastasis age, and parametrized by the expected age *a* when metastases become irremovable, treatment may remove metastasis. Also due to treatment, time to death is prolonged by a constant *h*.

The prognostic importance of cancer extent on patient outcome has been known for decades. The international standard for the staging of cancers is according to the the tumor/node/metastasis (TNM) system, published by the Union for International Cancer Control (UICC) for over 50 years, which classifies tumors based on local, regional and distant extension/spread (23). This classification is the most important determinant for patient management and prognostication. The TNM classification uses four stages to describe the extent of the primary tumor (T) based on the maximal diameter of the tumor (often measured in cm) and/or the infiltration depth in different anatomical structures; the definition of the stages is specific for every tumor entity. Given that larger tumors can be assumed to release more cells over time, they should have higher chances to develop metastases and therefore indicate a poorer prognosis. Studies analyzing the dependence of metastasis incidence on tumor size date back to the 1980’s (24). Importantly, patient survival was later observed to become independent of tumor size upon occurrence of metastases (25). This further supports the causative, and not only correlative, dependence of metastatic incidence on tumor size. The exact description, however, of how tumor size affects metastasis probability in the presence of colonization bottleneck and how patient outcome in turn depends on metastasis probability, is still lacking.

The metastatic process has been modeled using various mathematical approaches, based on different hypotheses about what affects metastatic probability and patient outcome. In their pioneering work, Liotta and colleagues proposed the first series of mathematical models of the metastatic process in mice (26–28). They also were the first to describe that larger implanted tumor clumps have a larger chance to seed metastases (21). That finding was not taken account for in theoretical models developed since that time. Assuming release of metastatic cells at a rate proportional to the volume of the colony raised to some power, models were proposed for the distribution of sizes of metastatic lesions in time (29, 30). Hartung et al. (31) modeled primary and metastatic growth in mice, assuming that metastatic cells are constantly released from the surface of the growing tumor with a given rate. Michor et al. (32) proposed a constant population size Moran process model of evolutionary emergence of pro-metastatic mutation in the primary tumor. This model was later extended to account for expanding tumor size and dependence on tumor size in the cell release probability (33), and consecutively put into a clinical context, showing a relation between tumor size, metastatic burden, and patient survival, accounting for surgery and chemotherapy (34, 35). This work allowed for calculation of probability of metastasis at a given time during tumor evolution, the expected number of metastatic sites, and the total number of cancer cells as well as metastasized cells. Haeno *et al*. (35) fitted parameters of this model to primary and metastatic cell count measurements determined post-mortem in pancreatic cancer patients. Benzekry *et al*. (36) modeled the non-linear dependence of the probability of metastatic relapse on primary tumor size, finding that between-patient variability of intrinsic metastatic potential is a key parameter, and suggesting that tumor size alone has a limited utility as a predictor of metastatic recurrence. None of the previous studies accounted for the metastatic bottleneck in a nontrivial way, other than considering very small export or pro-metastatic mutation rates, or their combination.

Unlike previous approaches, in the present work, in addition to tumor size, we explicitly account for the metastatic bottleneck in estimating metastatic probability. According to our model, the bottleneck in metastasis formation results from a stochastic process of metastatic colony initiation (phase 2) where the ratio of proliferation to death rates of cells in the colony increases with its size (Figure 1b). This assumption is motivated by experimental observations of (22, 26). In contrast to this assumption, once the metastatic colony grows large enough and reaches high enough density, the proliferation rate of its cells is naturally expected to decrease and the death rate is expected to increase due to overcrowding. In the case of metastasis initiation, however, the situation is the opposite, and cells in the growing colony should benefit from a more friendly environment once surrounded by more of their own kind. The proposed model quantifies the severity of metastatic bottleneck as the probability that the seeded colony survives. Based on this model, we find a critical colony size that separates very unlikely from very likely metastasis initiation (Figure 1c). To our knowledge, our model is the first to locate the rate limiting step of metastasis formation in the metastasis initiation phase. This is in contrast to the Haeno *et al*. model (34, 35), which implicitly identifies the rate limiting step with the emergence of pro-metastatic mutation in the primary tumor. Similarly to previous studies, we assume exponential primary tumor growth (34, 35), that tumors release metastatic cells from their surface (31), and that surgery removes the source of metastatic seeding (36). Based on these assumptions and on the model of metastatic bottleneck, we derive formulas for post-surgical metastasis detection and cancer death probabilities and any quantile (for example, median) time to death as functions of tumor diameter. Our formulas closely fit to and are predictive of epidemiological data for thirteen different cancer types. We correctly predict the median time to death for patients with clinically detectable metastases and the fact that it is independent of tumor size. Across cancer types, the median estimated probability of successful metastatic seeding is only of the order of 10^−9^, indicating that the initiation bottleneck is indeed the rate limiting step of metastasis formation. We estimate parameters characteristic of the cancer type aggressiveness: the expected time from the occurrence of metastases to patients death, the life prolongation achievable with treatment, the time for the metastases to grow large enough to become detectable, and the average age of metastases when they become irremovable by treatment (Figure 1d). We apply our model to predict the impact of different treatment decisions, such as surgery delay, increase of chemotherapy efficiency, and a potential therapy targeting the metastatic bottleneck. We conclude that the effect of such treatment decisions depends on the tumor size and the bottleneck severity and we predict that strengthening the metastatic bottleneck therapeutically would result in a dramatically stronger patient survival benefit as compared to chemotherapy. The presented theoretical results, predictions, and estimated parameters indicate that metastasis initiation is the rate limiting step of the entire metastasis formation process and therefore it should lay in the focus of medical research.

## 2. Results

### 2.1. Dependence of clinical outcome on tumor size as observed in epidemiological data

To comprehensively investigate how clinical outcome depends on tumor size, we systematically analyzed epidemiological data of fourteen cancer types from eleven primary sites, namely ductal and lobular breast, ovarian, endometrial, esophageal, gastric, colon and mucionous colon, rectal, pancreatic, non-small cell lung, head and neck, renal, and bladder cancer (Methods and Table S1). We selected a total of 159,191 patients from the Surveillance, Epidemiology, and End Results Program (SEER) database (37), only including patients with a single primary tumor that was surgically removed without prior treatment and where primary tumor diameter was measured (for details about patient selection see Methods). We investigated the dependence of metastasis detection frequency, cancer death frequency, and median time to cancer death on primary tumor diameter. For all cancer types except ovarian cancer, we found that metastasis detection frequency and cancer death frequency significantly increase with tumor diameter, while the median time to death decreases (Figure 2, Table S2). Ovarian cancer data may not follow these monotonic trends because ovarian carcinoma has a unique and very efficient way to rapidly spread within the peritoneal cavity, which differs markedly from the classical pattern of hematogenous dissemination (38). This deviating behavior further supports the connection between tumor size and metastatic probability via hematogenous tumor cell dissemination. Given that nearly all cancer deaths can be attributed to metastasis, cancer death frequency reflects the frequency of patients with metastases already present at diagnosis and not removed by treatment, including undetectable micrometastases. Accordingly, the frequency of metastasis detection at diagnosis is consistently smaller than cancer death frequency (Figure 2a). In agreement with previous observations made for breast cancer (25), median time to death does not depend on primary tumor diameter for patients with detectable metastasized tumors also for other cancer types that we analyzed. As expected, those patients die of cancer much earlier than patients without detected metastasis (Figure 2b).

**Figure 2.**
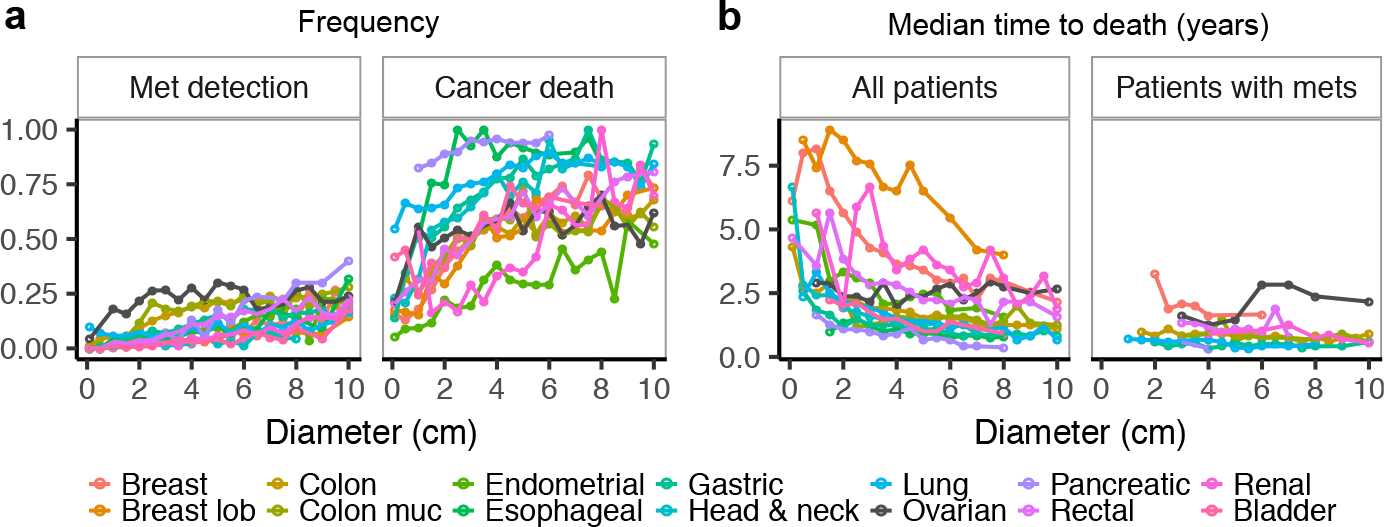
Epidemiological data from the SEER database (37) for different cancer types (colors) **a** The probability of metastasis detection (left) and the probability of cancer death (right) increase with tumor size at diagnosis. Only ovarian cancer data (black) do not follow these monotonic trends. **b** Median time to death for all patients (left) decreases with tumor size at diagnosis, again except for ovarian cancer (black). For the subset of patients who had metastases detected at diagnosis (right), the median time to death is generally much shorter and is not tumor-size dependent.

### 2.2. Mathematical model of metastatic bottleneck, its impact on metastasis probability and on clinical outcome

To describe the first and efficient phase of metastasis formation, we assume that as the primary tumor grows, it continues to release cells from its surface. Primary tumor growth is modeled as an exponentially increasing spherical volume with the doubling time set to constants measured for different cancers (Table S3). The per-cell per-year release rate is fixed to match experimental data, showing that a tumor of one gram contains about 10^9^ cells (39) and that such a tumor sheds around 1.5 × 10^5^ cells per day (26). The extravasation probability is fixed to 0.8, based on experimental observations in mice (15) (Methods).

To model the second, extremely inefficient phase of metastasis initiation, we rely on the observation that metastatic propensity increases with colony size (21, 22), which is expected for colony outgrowth up to certain limit size. Accordingly, our model is an inhomogeneous birth-death process, where the ratio of proliferation to death rates of each individual cell depends on the colony size with a proportionality constant b, termed the bottleneck severity (Figure 1b). From this assumption, we derive the probability of successful metastatic seeding starting from a single cell, *s*_1_(*b*) = *e*^−*b*^ (Methods). The bottleneck severity is the critical number of cells, where, as the colony size increases from smaller than b to larger than *b*, the probability of survival of the colony grows dramatically from very small to very large (Figure 1c). Up to the time when it reaches diameter *d*, the primary tumor releases a very large number *N*(*d*) of cells that can be regarded as independent trials to initiate metastases, each with very small success probability *s*_1_(*b*). For a patient with tumor size *d*, the number of metastases is thus Poisson distributed with rate *N*(*d*) · *s*_1_(*b*), and the probability of having at least one successfully initiated metastasis is 
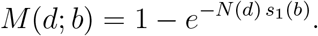

Importantly, the bottleneck for an individual metastatic colony and for an individual patient may be different, depending on the genetic makeup of the tumor cells, the organ where they try to seed, and due to differences in the immune system of the patients. We model this variability of the bottleneck severity by assuming that it is log-normally distributed with parameters *μ* and *σ*. Also, we assume that complete removal of primary tumor by surgery eliminates the source of metastatic seeding. To model post-surgical patient outcome, we make the following assumptions (Figure 1d). Once metastasis initiation succeeds in one site of the body, it takes an average time *δ*_0_ for the metastasis to become lethal. Time *δ*_1_ is required for the metastases to grow large enough to be detectable at diagnosis. Removal of metastases by systemic therapy is less probable as they grow, and the time when the metastases become irremovable is exponentially distributed with expectation *a*. Finally, we consider that treatment prolongs survival of patients on average by h years. The six quantities *δ*_0_, *δ*_1_, *a*, *h*, *μ*, and *σ* differ for each cancer type. They are the free parameters of our analytical formulas for *i*) probability of cancer death and for *ii*) probability of metastasis detection, as well as for *iii*) quantile time to cancer death for all patients and for *iv*) quantile time to cancer death for the subset of patients with detected metastases (Methods).

### 2.3. Model fitting and validation

To estimate model parameters for each cancer type, we fitted the formulas for metastasis detection (Equation 13) and quantile times to death (Equation 15) for all patients and eleven different quantiles to the SEER data (Methods). These formulas together depend on all six free parameters of our model. Although the model can predict the frequency of cancer death and median survival for the subset of patients with detected metastases, this data was not used for model fitting but kept for validation. The theoretical model curves fit the data extremely well (Figure 3 and Figure S1, blue curves). Fits to other quantile time to death data are comparable to the fits to the median (Figure S2). The ovarian cancer data was excluded from this analysis because it was the only cancer type which did not show dependence of patient outcome on tumor size, and therefore did not fit our model assumptions (Figure 2).

**Figure 3.**
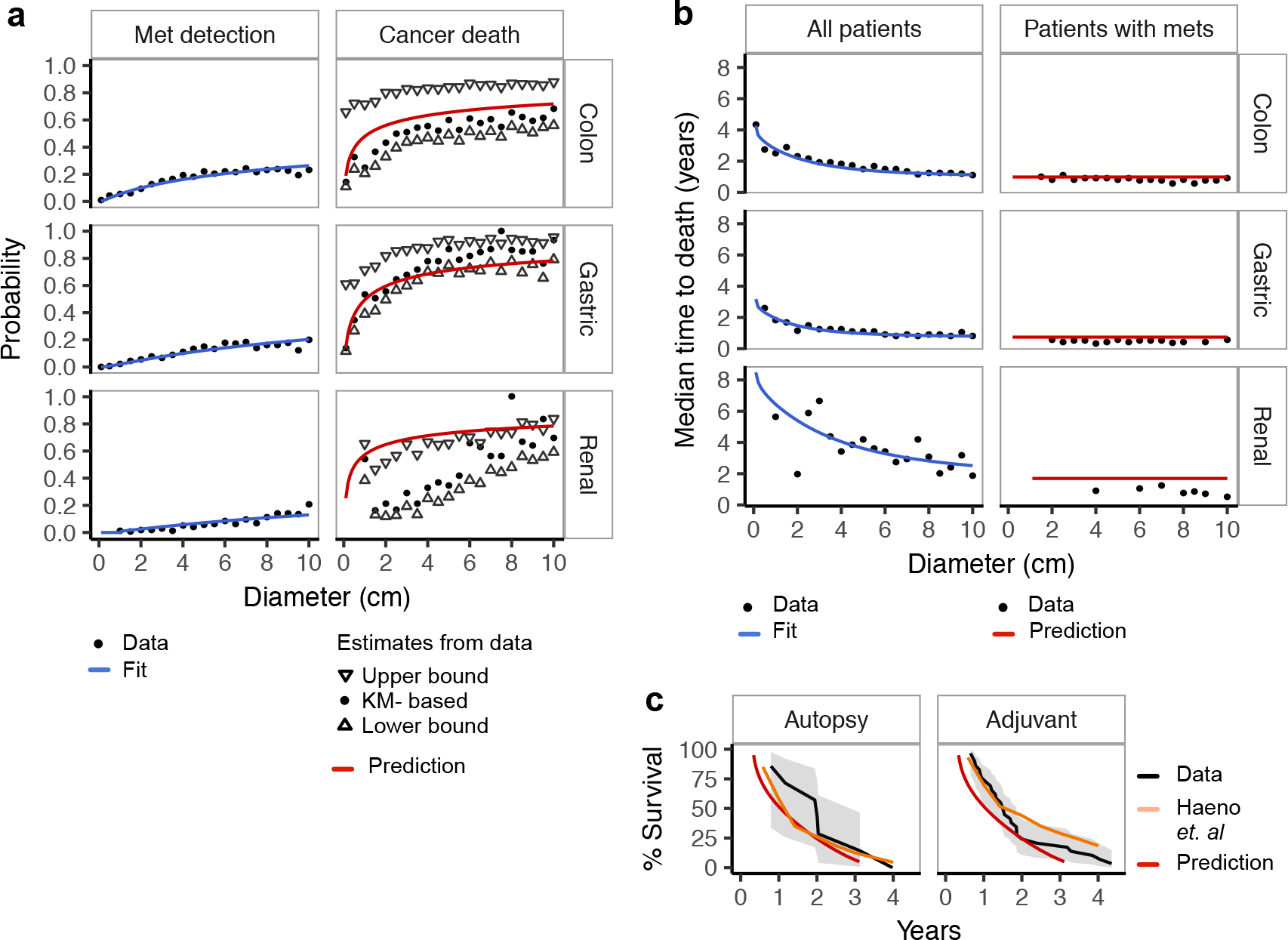
Model fit and validation. **a,b** Fit and validation of our model on the SEER data, exemplary for three cancers, for which we obtained the best, intermediate, and worse fit, respectively (see Figure S1 for remaining ten). Black points represent data records, blue lines show fitted, and red lines predicted curves. **a** For cancer death probability, up and down-oriented triangles present the upper and lower estimates of that variable from the data, respectively, while the black dots represent a Kaplan-Meier (KM) derived estimate (Methods). In agreement with the data, we model cancer death frequency (right) as the probability of metastasis that were not removed by treatment, which accounts also for undetectable metastases and is thus larger than metastasis detection probability (left). **b** Despite the fact that the median time to death data for all patients (left panel), which was used for model fitting, is tumor-size dependent, the model correctly predicts the much shorter median time to death for patients with metastases detected at diagnosis, and that this time is virtually constant across tumor diameters (right). **c** Independent validation on survival data for two pancreatic cancer cohorts: Autopsy and Adjuvant, analyzed by (35) (black, with gray confidence bands). We predict their survival function using our pancreas cancer model, fit solely to SEER data, and accessing only information about tumor diameters of patients in the cohorts. For both cohorts, our predictions (red curves) are as close to the data as predictions obtained from the the Haeno *et al*. model (orange).

With all parameters estimated, we next validated the model by comparing the model predictions of cancer death probability (Equation 12) and quantile time to death for patients in which metastases were detected at the time of diagnosis (Equation 17) to the corresponding data in the SEER database. Due to deaths for other reasons than cancer present in the SEER records, it is hard to evaluate the frequency of cancer death from the data. We thus calculated three estimators of that variable. The upper bound estimator treats all other deaths as cancer death, the lower bound estimator treats all patients died from other reasons as alive, and the third estimator is Kaplan-Meier derived, treating other deaths as censoring (Methods). We find that the model predictions prove consistent with cancer death frequency data, such that for most cancer types our predictions are very close to one of its estimators (Figure 3a and Figure S1a, red curves). Remarkably, although the quantile time to death for all patients used for fitting decreases with tumor diameter, the model very closely predicts the essentially constant and much lower level of the median time to cancer death for patients with detected metastases (Figure 3b and Figure S1b, red curves).

Additionally, we validated our model on independent survival data from two pancreatic cancer cohorts, named Autopsy and Adjuvant (35). Using our formulas that we fitted to pancreatic cancer data from the SEER, we very closely predict the survival functions of these two independent cohorts using only their primary tumor diameters (Figure 3c; Methods). Our predictions are as close to the data as those obtained from the model of (35), which, however, was fitted to the Autopsy cohort itself, having access not only to tumor diameters and clinical information but also to counts and sizes of metastatic lesions determined at autopsy.

### 2.4. Characterization of cancer types with model parameter values

Based on the estimated model parameters, we ranked cancer types in terms of their severity and susceptibility to treatment. The estimated time intervals from the formation of a successful metastatic colony until the patient’s death, *δ*_0_, are different for different cancer types and span 3 to 12.6 years (Figure 4a). We infer that the times *δ*_1_, during which the metastases grow detectable, are very close to *δ*_0_. This indicates that once a metastasis is clinically observable, the disease is already at a very late stage. This is why our model predicts that the median time to death is determined by treatment induced life prolongation h and becomes independent of the primary tumor size (Figure 4b; compare Figure 3b). For all cancer types, the estimated prolongation h is much shorter than the intervals *δ*_0_ and *δ*_1_, with the largest values of around 1.7 years. The times for the metastases to become lethal (*δ*_0_) and detectable (*δ*_1_), as well as life prolongation *h* tend to be shorter for cancer types that are known to be more aggressive, such as pancreatic or esophageal carcinomas. The expected age when the metastases become irremovable *a* was either estimated around 1 year (its upper bound; Methods) or only several weeks (Figure 4c). Importantly, the estimated distribution of the bottleneck severity *b* provides new insights into the number of tumor cells critical for successful metastasis initiation. The distribution is similar across all analyzed cancers, with one outlier of colon mucinous cancer, which has both the largest mean and variance (Figure 4d). The most common median bottleneck is 19 cells, corresponding to a metastatic success probability of *s*_1_ = 5.6 × 10^−9^. Such small probabilities of successful metastasis initiation inferred across different cancer types agree with the known inefficiency of this phase of metastasis formation, arguing in favor of our bottleneck model. For all cancers, the bottleneck distribution has a large variance, indicating considerable variability of bottleneck severity within and between patients.

**Figure 4.**
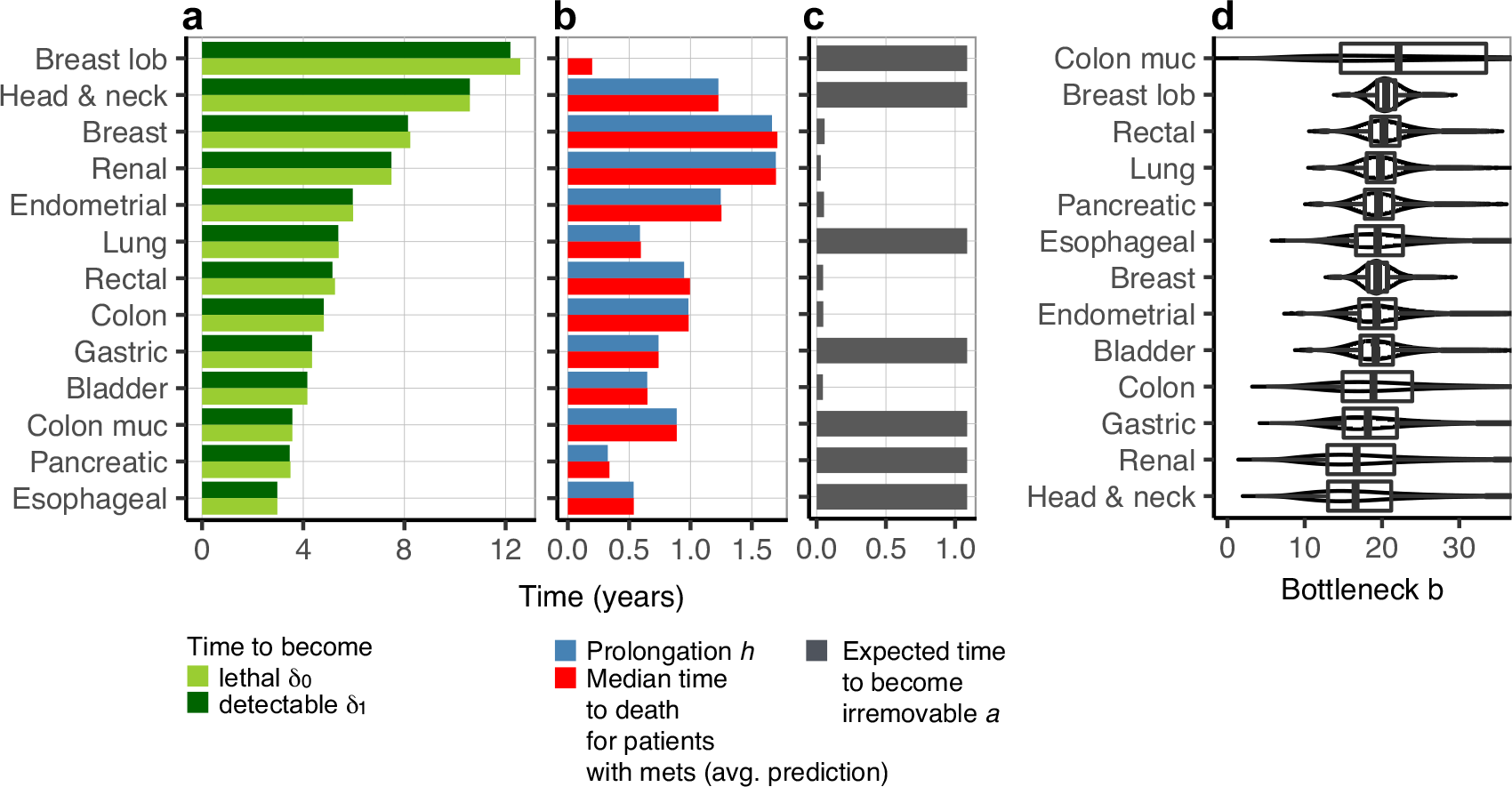
Estimated model parameters for thirteen cancers. **a** The estimated time for metastases to become lethal, *δ*_0_ (light green) and the time for metastases to become detectable, *δ*_1_ (dark green), are shorter for more aggressive cancers (**a**). **b** The estimated life prolongation due to treatment *h* (blue) is very close to the predicted median time to death for patients with metastases detected at diagnosis, averaged over tumor diameters (red; compare Figure 3b). **c** The expected time of metastases to become irremovable by treatment, a, was estimated to be the upper bound of around 1 year, for seven cancers. For the remaining six cancers, *a* is on the order of weeks and the removal probability decreases much faster as a function of metastasis age. **d** Box plots showing 25th, 50th (median), and 75th percentiles (vertical bars) and 1.5 interquartile ranges (horizontal line ends) of the bottleneck severity *b* (x-axis) for all cancers, ordered by the median severity (y-axis). The median bottleneck severities range from 17 to 22, with the most common median severity being 19 cells. Large variances of the bottleneck distributions indicate strong variability in bottleneck severities both within and between patients.

### 2.5. Predicted impact is most dramatic for treatment aiming at increasing bottleneck severity

Finally, we apply our model to predict impact of three different treatment decisions on cancer death risk (Figure 5 and Figure S3). Specifically, we consider how cancer death probability will increase or decrease due to (*i*) delay of surgery by 16 weeks, modeled by effectively allowing the tumor to grow and seed the metastases 16 weeks longer than the actual time of diagnosis (Figure 5a), (*ii*) applying a more efficient chemotherapy, where the expected age at which a metastasis becomes irremovable is increased by 20% (Figure 5b), and (*iii*) a treatment strategy resulting in 20% increase of the median bottleneck severity (Figure 5c; Methods). Our model predicts that the impact of all three treatment decisions is tumor size-dependent, with lower impact for small and large tumors and larger impact for mid-sized tumors. Similar behavior, but only for the impact of surgery delay, was reported in mice by (36). Intuitively, decisions such as delay of surgery or modification of treatment efficacy may not matter both for small tumors, which did not yet succeed to develop metastases, and for large tumors, which already have acquired them. As the size of the primary tumor grows from small to mid-size, the probability of having metastases increases, and thus delay or additional treatment might have the greatest effect. Importantly, different cancer types differ in the primary tumor sizes that are most sensitive for treatment decisions.

**Figure 5.**
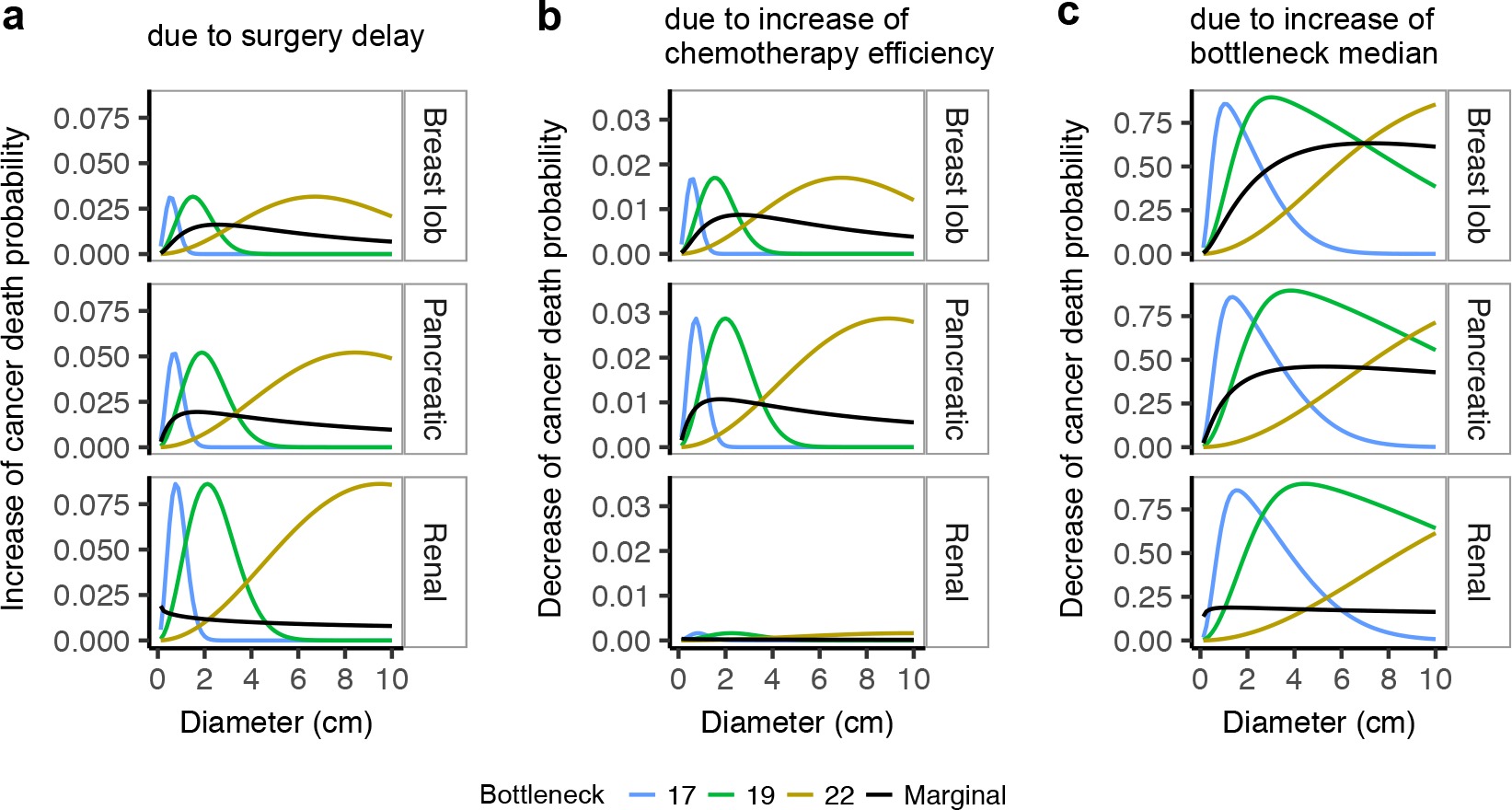
Predicted impact of treatment decisions. Change in cancer death probability (*y*-axis) due to treatment change depends on tumor diameter (*x*-axis) and bottleneck severity (colors), shown for three most impacted cancers (rows). Black curves present the change in cancer death probability, marginalized over the bottleneck severity. The increase of cancer death probability due to surgery delay by 16 weeks (**a**), as well as decrease of cancer death probability due to increase of chemotherapy efficacy by 20% (**b**) are both much smaller than the decrease due to strengthening of the metastatic bottleneck by 20% (**c**).

Besides the primary tumor size, our model suggests that the impact of treatment strategies depends also on the metastatic bottleneck, such that the peak of the impact is different for different bottleneck severities. For example, for fixed bottleneck severities (color curves in Figure 5a), the largest predicted increase of cancer death probability due to surgery delay of 16 weeks is for renal cancer. For a bottleneck severity of 19 cells and a tumor diameter of 2 cm, the increase is circa 0.9 and decays to 0 for larger tumors. For the same cancer type, but bottleneck severity 17, the peak impact of surgery delay is at a tumor diameter of 1 cm. The marginal increase of cancer death probability due to surgery delay of 16 weeks (black curves in Figure 5a) does not exceed 0.02, and is the largest for pancreatic cancer and a tumor diameter of 1.6 cm. The estimated bottleneck distributions (Figure 4d) imply that the impact will vary both between and within patients, depending, for example, on their immune system and affected organs. Since the individual bottleneck severities are unknown, our estimates of the marginal impact can be used to indicate how treatment decisions will affect most of the individuals.

Importantly, we predict that a systemic therapy that would decrease the chances of successful metastatic colony initiation and tighten the median bottleneck by 20% (Figure 5c), would roughly have a 100-fold higher impact on patient survival than a 20% increase of chemotherapy efficacy (Figure 5b). Across all cancers and diameters, the largest obtainable marginal decrease in cancer death probability due to increase in chemotherapy efficiency is just 0.01 (for pancreatic cancer and tumor diameter 1.7 cm). For a median bottleneck severity increase by 20%, the maximum marginal decrease of cancer death probability is 0.63, for breast lobular cancer. Recent advances in immunological cancer treatment led to clinical trials of anticancer vaccines (40), and a bottleneck shift may be achieved in the future, for example, by vaccines that strengthen the immune system against forming metastatic colonies, or by drugs that reduce cell adhesion in a cancer cell-specific manner.

## 3. Discussion

The presented work extends established knowledge about metastasis formation in several ways. Most importantly, although the extreme inefficiency of metastatic colonization is commonly accepted, to our knowledge, this is the first mathematical model that explicitly takes it into account and provides an explanation for the phenomenon of metastatic bottleneck. Inspired by experimental observations (21, 22), the main assumption of the model is that the ratio of proliferation to death rates increases with colony size. This assumption implies that, when passing a certain critical size, the probability that the colony will manage to survive switches rapidly from extremely low to very high. Furthermore, our work takes the effort to quantify the severity of this tremendous bottleneck from epidemiological data for different tumor types. To this end, for each cancer type, we estimate the distribution of the critical metastatic colony size and of the probability that the seeded colonies will survive. Third, it is also well known that tumor size influences prognosis and tumor diameter is one important factor composing the TNM staging system. We specify this association by linking it with metastatic probability and provide mathematical formulas that describe the non-linear dependence of several different patient outcome variables on tumor size. Importantly, from our model we deduce that the tumor size is also predictive of how much a patient will benefit from a given treatment. Model predictions from epidemiological data show that patients with larger or smaller tumor benefit less from additional treatments. This has important implications for clinical study design and would argue that exact tumor size should be measured and included in data analysis. In addition, our model suggests that, apart from tumor size, also the individual bottleneck severity has a significant impact on patient outcome. The model predicts a high potential of treatment aiming at strengthening the metastatic bottleneck, out-competing what can be achieved by standard therapies simply reducing tumor cell burden. Intuitively, such treatment would increase the chances of patient survival by reducing the chances to form a metastasis, for example by boosting the immune response. Finally, it was reported for breast cancer already before, that the time to death for patients with metas-tases detected at diagnosis becomes independent of tumor size (25). Our analysis of SEER data demonstrates, and our model correctly predicts, the same phenomenon also for other cancer types.

Several modeling choices have been made in the featured analysis. For instance, similarly to Haeno *et al*. (34, 35), we assume deterministic exponential growth of the tumor volume (41). Gompertzian growth is considered more suitable to describe the initial, exponential-like increase of the tumor volume and its later leveling off for larger tumors. The key factor in our model, however, is the probability of acquiring the first metastasis, which is most likely to happen for smaller than for larger tumors, where the exponential tumor growth is adequate. Second, contrary to (34, 35), we do not consider emergence of pro-metastatic mutations, but rather metastatic colony initiation as the rate limiting step of metastasis formation. Likely, both are independent factors contributing to the extreme inefficiency of this process. Although we believe and literature (4–7, 15, 18) supports that metastasis initiation is the main factor, this needs to be determined in future experimental studies. Determination of the rate limiting factor is crucial for clinical handling of metastatic tumor disease, as the implications for therapy are entirely different depending on whether it is the emergence of mutations or the initiation of metastatic colonies. The former case is out of therapeutic access, as we can do essentially nothing to prevent random mutations. On the contrary, in the latter case, there is a potential that the bottleneck is controllable by therapeutic means, such as immunotherapies, once better understood on a biological level. Perfect agreement of the proposed model with the data it was fit to, and more importantly, its predictive power validated on unseen data, indicate that our model captures the essence of the mechanisms behind metastasis formation, and that the tumor size, metastatic initiation bottleneck, and the probability of metastasis occurrence are together the key to describe clinical outcome such as patient survival or risk of cancer death.

We deliberately kept the model as simple as possible, minimizing the number of free parameters and model variance. Except from modeling the distribution of the bottleneck severity, which is the most sensitive parameter in the model, other parameters are introduced as robust summary statistics of time to events, rather than their distributions. For example, the model depends on the average time to death from the acquisition of metastases. Similarly to parameters, the modeled clinical variables are also robust statistics, such as quantile time to death. Consequently our model is less expressive than other models, describing for example metastasis size distribution (29, 30). As such, it can be fit to and be predictive of epidemiological data, while more expressive models require model-tailored meticulous measurements of tumor burden. Despite its simplicity, our model is able to capture and be predictive of a surprising variety of aspects of clinical outcome. Although model parameters were fit to one type of data, namely metastasis detection and quantile time to death as functions of tumor volume, it was able to closely predict different variables in the data that were unseen during model fitting, namely cancer death probability and the median time to death for the subset of patients with metastases detected at diagnosis, as well as independent survival data of (35). In summary, the presented model is an important step forward in the understanding of metastasis formation, its bottleneck, and its impact on patient outcome.

## Acknowledgements

We thank the Center for Interdisciplinary Research (ZiF), Bielefeld University, for partly funding and inspiring this work, via involvement of all authors in the ZiF Cooperation Group “Multiscale Modeling of Tumor Initiation, Growth and Progression: From Gene Regulation to Evolutionary Dynamics” from September to December, 2016. E.S. is funded by 2015/19/P/NZ2/03780 grant from the National Science Centre, Poland. T.K. is grateful to his Department of Control Systems and Mechatronics, Faculty of Electronics, Wroclaw University of Technology for funding several visits to Dresden, Warsaw and Basel.

## Author contributions

E.S., T.K. and N.B. designed the mathematical model. E.S. acquired and analyzed data and wrote the manuscript with critical comments from all the authors. B.K. contributed to data acquisition and analysis. E.S., T.K., B.K. and N.B. contributed to study conception and design, interpreted the data and provided a critical reading of the manuscript.

## Methods

### Processing epidemiological data

Patient data was downloaded from the SEER database (37), with last follow-up date December 31, 2013. We identified 2,091,631 tumor records with given SEER histology and behavior codes that corresponded to one of fourteen common cancer types. From this, we selected the 549,835 records for adult patients with exactly one tumor of precisely coded diameter below 10 cm at diagnosis, who had surgery but no prior radiation treatment. Tumors were restricted to below 10 cm since there were not enough records for tumors above 10 cm for reliable estimation of modeled variables. To estimate quantile time to death correctly, the patients follow up had to be long enough to result in a representative data distribution. Thus, we defined minimum follow-up times for every cancer type (Table S1) leading to a selected cohort of 159,191 patients that fulfilled all criteria. From this cohort, different subsets of patient records were defined to estimate different quantities.

To estimate cancer death frequency, we only used records for patients for whom their survival time was recorded, leaving 110,102 records. We grouped these patients by cancer type and tumor diameter on a grid from 0.5 to 10 cm, filtering out groups with less than twenty patients to avoid fluctuations due to small sample sizes. For each group, we aimed at assessing the ratio of patients who died of cancer to the total number of patients in the group. Here, we had to account for the fact that some patients die of reasons other than the tumor disease, and therefore we cannot determine whether they would eventually die of or survive cancer. We thus calculated three estimators of cancer death frequency. For the upper bound estimator, all other deaths are treated as deaths from cancer, while for the lower bound, all dying of other reasons are treated as survivals. The intermediate estimator is computed as 1 minus the Kaplan-Meier estimate of the survival function at its last measured time point, where the survival times of both the other deaths and alive patients contribute to the censored cases.

From the pool of 159,191 patients, metastasis detection frequency was assessed for the subset of 158,358 who had metastasis examined. To this end, for each cancer and tumor diameter group of at least 20 patients, we calculated the fraction of the group who had distant metastases detected at diagnosis.

To extract quantile times to death for all patients, we limited the pre-selected 159,191 patients to 47,166 whose survival time was recorded and who were reported to have died of cancer. For each group of at least twenty patients per each cancer type and each diameter, we calculated their quantile times to death for eleven different quantiles, from the 0.35-th to 0.65-th quantile. For example, the 0.5-th quantile time represents the median time to death, i.e., the number of years for which half of the patient in a group had died of cancer earlier than this number of years.

Finally, we studied a much smaller subset of 9,159 patients, who had metastases examined and the metastases were detected at diagnosis. Again, we evaluated their quantile time to death in cancer-and diameter-resolved groups of at least twenty patients. Here, with the low number of available records, the patient groups were large enough to generate at least two data points for only eight of thirteen cancer types.

Since the SEER database provides only the time of diagnosis, we assumed that diagnosis, surgery, screen for metastatic sites, and tumor measurement took place at the same time point.

### Metastasis initiation model

The model describes the formation of a metastatic colony starting from a single tumor cell that had infiltrated into a secondary organ. We assume that initially, the larger the colony manages to grow, the easier it is for its cells to divide. At this stage, cell population is much too small for the tumor cells to compete against each other for nutrition and oxygen, and to exhibit hypoxia and necrosis, which result from over-crowding. Thus, we model the initial metastatic colony growth as a continuous-time binary branching process, where each state *i* is the total number of cells in the colony. We define the ratio of the cellular proliferation rate λ_*i*_ and the death rate *μ*_*i*_ as an increasing function of *i*, 
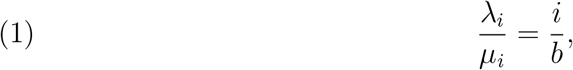
where *b* is a proportionality constant, which we call the bottleneck severity. Intuitively, the larger the severity *b*, the stronger the metastatic bottleneck. Indeed, the larger *b*, the smaller the proliferation to death rate ratio. For a population of *i* cells, the individual rates λ_*i*_ and *μ*_*i*_ correspond to the population proliferation rate *i* · λ_*i*_ = *i*^2^ *μ*_*i*_/*b* and the population death rate *i* · *μ*_*i*_, respectively.

Let 
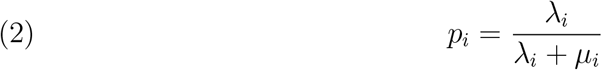
denote the jump probability from state *i* to *i* + 1, and 1 − *p*_*i*_ the probability for the jump from *i* into *i* − 1. The jump probability fully describes the stochastic dynamics of the population, and it depends only on the combined proliferation and death rates. Intuitively, for large jump probability *p*_*i*_ the colony size will likely increase in the next step, while for small *p*_*i*_ it will likely decrease. The critical colony size is reached when this probability passes 1/2. More precisely, the smallest total number of cells that results in jump probability larger or equal than 1/2 is called the critical size. Inserting the rates in relation described by (1) into Equation (2), we find that the critical size is equal to ⌈*b*⌉.

This branching process is a birth and death chain with a single absorbing state *i* = 0, corresponding to colony extinction. We refer to the probability *s*_1_ of never reaching the absorbing state starting from a single initial cell as the metastasis success probability, given by (Supplementary Methods) 
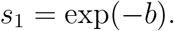

Hence, the metastasis success probability exponentially approaches 0 with rate equal to the bottleneck severity *b*. To make the dependence of *s*_1_ on the bottleneck severity *b* explicit, in further considerations we will use the notation *s*_1_(*b*).

The probability of colony survival (not going extinct), starting from any number of cells *i* > 1 (Supplementary Methods) reads 
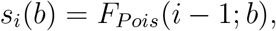
where *F*_*Pois*_(*i*; *b*) denotes a cumulative Poisson distribution function with parameter *b*. Thus, as the colony size passes the critical number of *i* = ⌈*b*⌉ cells, the probability of survival from *i* cells *s*_*i*_(*b*) passes 0.5, rapidly switching from small to large values (Figure 1c).

Three aspects of the presented model are discussed more deeply in Supplementary Methods. First, it is of course not possible that the proliferation to death ratio goes to infinity with increasing colony size. We show, however, that a model assuming a truncation to a constant ratio after the colony grows large, would yield only negligibly different probabilities of colony survival. Second, we discuss that the model described here does not specify the timing of the stochastic process of colony initialization, but that it can once average cell death rate was measured. Finally, here we simply assumed that the dependence of proliferation to death ratio on *i* is linear. Theoretical extension of the model to a more general case, where 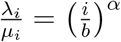, with addition of parameter *α*, is introduced.

### Metastasis probability for patients with given tumor size

Denote *t*(*d*) the time, measured in years, that elapsed from the onset of the tumor to the time of diagnosis with tumor diameter *d*. Let *N* (*t*), for *t* ≤ *t*(*d*), be the total number of cells that were released from the primary tumor up to time t and managed to extravasate to the secondary organ. The value of *N* (*t*) depends on the primary tumor growth, per-cell per-year release rate, and extravasation probability. We regard the fate of each released tumor cell as independent. *N* (*t*) is interpreted as the number of independent trials that the tumor cells made in order to seed metastases, while *s*_1_(*b*), defined by the metastasis initiation model, as their success probability. Given that *N* (*t*) is very large, while the metastasis success probability *s*_1_(*b*) is very small, we approximate the distribution of the number of metastatic sites at time t by a Poisson distribution with expectation *s*_1_(*b*) · *N* (*t*). The metastasisprobability *M*(*t*; *b*) for time *t* is then defined as the probability of developing at least one metastatic site up to time *t*, equal to one minus the probability of having zero metastases under the Poisson distribution 
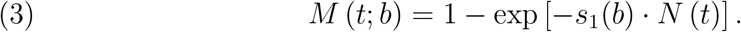

Importantly, *M*(*t*; *b*) defines also the cumulative waiting time distribution for the first successful metastasis. Let *τ* be the random variable describing the waiting time from the onset of the primary tumor to the first metastasis creation. Indeed, the cumulative distribution function for *τ* equals 1 minus the probability that no metastases have occurred up to time *t* 
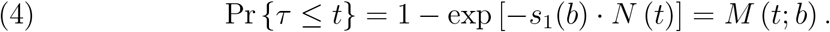

For a given fixed time point *t*\* ≤ *t*(*d*), to compute *N* (*t*\*), we consider the per tumor instantaneous rate *Y*(*t*) of metastatic cell spread, such that 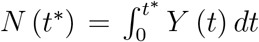. The rate *Y*(*t*) increases with the tumor volume *X* (*t*), measured in cm^3^. With additional assumptions of spherical tumor shape and that the majority of cells released from the tumor originate from the tumor surface, 
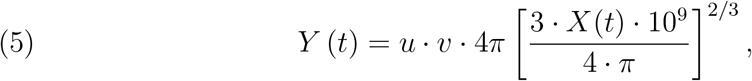
where the constant *u* is the per-cell per-year release rate of cells from tumor surface, *υ* is the extravasation probability, *X*(*t*) is the volume of the tumor in cm^3^ and 10^9^ gives the number of cells in unit (1cm^3^) volume. To set a realistic release rate *u*, we assumed that for all cancer types, there are 10^9^ cells in a tumor weighting a gram (39), and that around 1.5 × 10^5^ cells are shed from such tumor per day (26), corresponding to around 547.5 × 10^5^ per year. Given that such a tumor of spherical shape has around 4.836 × 10^6^ cells on its surface, we obtained that the per-cell per-year release rate u can be fixed to 11. The extravasation probability v is fixed to 0.8, based on experiments following the fate of tumor cells injected into mice, which stated that around 80% of cells managed to extravasate to secondary organs within three days (15).

To model the tumor volume as a function of time we assume a basic deterministic model of exponential tumor growth (41) 
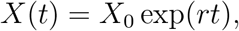
where at time *t* = 0 the tumor volume is *X*_0_ = 10^−4^cm^3^, corresponding to 10^5^ cells (with volume of a single cell of 10^−9^cm^3^). This initial condition *X*_0_ is chosen to make sure that with *X*_0_, the tumor growth has already passed its early stage of low cell count, in which growth dynamics may be stochastic rather than deterministic. The growth rates *r* are different for each cancer type and fixed to literature-derived constants (Table S3). With these assumptions, we can compute the integral 
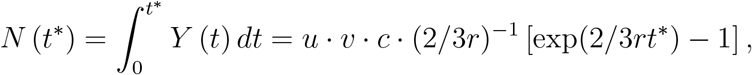
where the constant 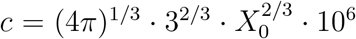. This fully defines *M*(*t*; *b*) (Equation 3).

With the assumptions of exponential growth for the primary tumor and from the volume formula for its spherical shape we have 
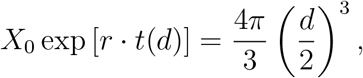
and obtain that a tumor of diameter *d* has age 
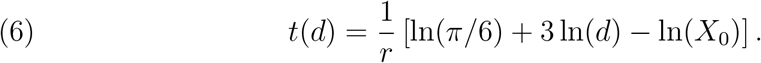

To define the metastasis probability as a function of diameter instead of time, we simply use Equation (4) together with (6) and denote 
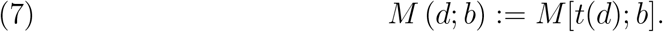

Importantly, metastasis probability *M* (*d*, *b*) is expected to be much larger than the metastasis detection probability, i.e., incidence of detected metastases in patients with the same tumor size, as it accounts for the probability of having micrometas-tases, which are not detectable by current screening techniques.

### Cancer death probability for patients with given tumor size

Given the fact that metastases account for circa 90% of cancer deaths, the cancer death probability should depend on the metastasis probability. To make this relationship explicit in the model, we first assume that in order to die of cancer, it is sufficient to have at least one successful metastasis at a distant site. Second, we focus only on patients who had tumors resected following the diagnosis, and assume that the surgery fully removes the source of metastatic seeding. Third, we also account for the trivial fact that patients presenting at diagnosis are alive and their lives were thus not yet threatened by the metastases. To this end, we assume that there is a certain, cancer-specific time *δ*_0_ that on average takes the metastasis to eventually become lethal in the case when there is no treatment. The metastasis probability becomes a conditional probability, denoted *M*(*t*; *b*, *δ*_0_), conditioned on the fact that up to time point *t*_0_(*d*) = *t*(*d*) − *δ*_0_, where *t*(*d*) is the time of diagnosis, no successful metastasis has been created. This conditional probability can be computed as 
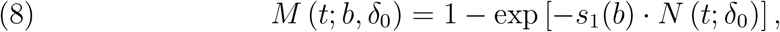
where for a given time point *t*\* 
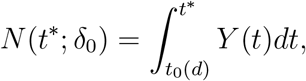
and *Y*(*t*) is the instantaneous per tumor rate of cell spread, given by Equation 5. Computing the integral with the assumption of exponential tumor growth we obtain 
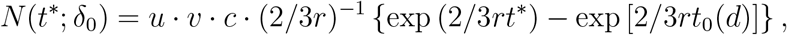
where the constants *u*, *υ*, and *c* are defined as above. The conditional metastasis probability as a function of tumor diameter is defined, similarly to Equation 7, by *M*(*d*; *b*, *δ*_0_) := *M*[*t*(*d*); *b*, *δ*_0_]. Fourth, we incorporate effects of treatment in the model, accounting for the fact that post-surgical therapy, most commonly chemotherapy, may eradicate some of metastases that could be life-threatening without treatment. We assume that the metastasis removal is less likely for older, and thus larger metastases, and that the time *t* ≥ 0 for the metastases to become irremovable is exponentially distributed with *a* cancer-specific parameter 1/*a*, where a corresponds to the expected age of metastases when they become irremovable.

Thus, for metastases which originated *t*′ years before the diagnosis, their probability of being removed be treatment is the probability that they became irremovable at some time point *t* > *t*′ 
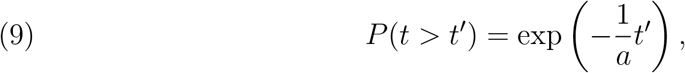
and decreases with its age *t*′.

For a given bottleneck severity *b*, we model cancer death probability as a function of time from tumor onset *t* as the post-treatment, conditional metastasis probability *M*(*t*; *a*,*b*,*δ*_0_) that at least one metastasis was successfully initiated by the time t and was not later removed by treatment 
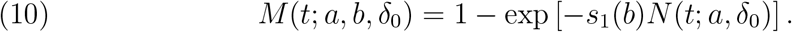

Here, for a given time point *t*\* 
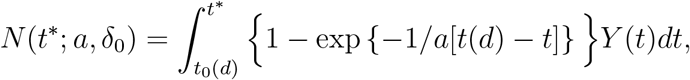
where *t*(*d*) denotes the time of diagnosis with tumor size *d*, and the instantaneous per tumor cell spread rate *Y*(*t*) (Equation 5) is multiplied by the probability that metastasis, which is *t*(*d*) − *t* years old, is not removed by treatment after the diagnosis. Solving the integral, we obtain 
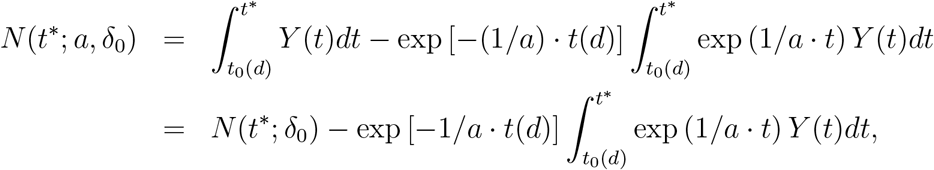
where the integral in the second term can be further simplified to 
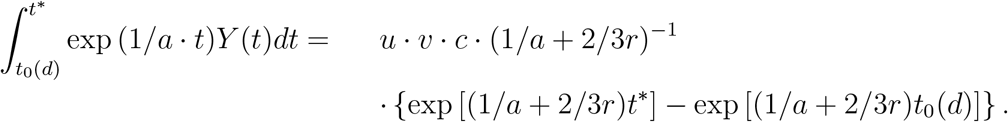

Finally, we model the variability of the bottleneck severity *b*. Here, we follow the principle, that the bottleneck severity, which determines the tremendously small metastatic success probabilities *s*_1_(*b*) = exp(—*b*) and is thus an extremely sensitive parameter, should be modeled with a probability distribution, and not just by the mean value, as we do with other parameters. Bottleneck variability is also expected due to biological reasons, such as differences in immune system strength and metabolic rate between patients, in genetic makeup and metastatic potential between tumor cells, as well as in the different organs within the body. The actual values of bottleneck severity *b* are unknown and with current technology cannot be measured. We assume that *b* is a random variable from a log normal distribution with density *f*(*b*; *μ*, *σ*), and cancer-specific parameters location *μ* and scale *σ*.

Taking all these considerations together, for patients who underwent tumor surgery followed with therapy reducing metastatic load, their cancer death probability as a function of time from tumor onset is given by the post-treatment, conditional metastasis probability marginalized with respect to the distribution of *b* 
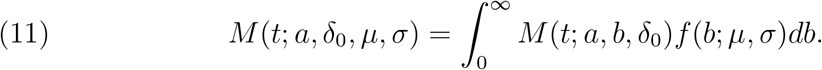

Cancer death probability as a function of tumor diameter *d* becomes 
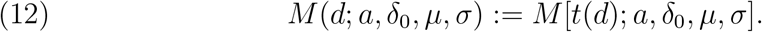

Thus, the formula for cancer death probability depends on the set of cancer-specific parameters {*a*,*δ*_0_,*μ*,*σ*}.

### Metastasis detection probability in patients with given tumor size

To model metastasis detection probability we make similar assumptions as above, namely that patients presenting with a tumor of size d should not have developed metastases before *t*_0_(*d*) = *t*(*d*) − *δ*_0_, where *t*(*d*) is the time of diagnosis and *δ*_0_ is the cancer-specific parameter corresponding to the average time for the metastasis to become lethal. We assume that each metastasis needs a mean time *δ*_1_ till it reaches a detectable size. With these assumptions, for a given bottleneck severity *b*, metastasis detection probability depends on the conditional metastasis probability *M*[*t*_1_(*d*); *b*, *δ*_0_] (Equation 8), that at least one successful metastases was created within the time interval [*t*_0_(*d*); *t*_1_(*d*)], where *t*_1_(*d*) = *t*(*d*) − *δ*_1_ and *δ*_1_ < *δ*_0_. Finally, we account for the bottleneck variability by assuming it is log normal distributed with parameters *μ* and *σ*, and obtain metastasis detection probability as a function of tumor diameter 
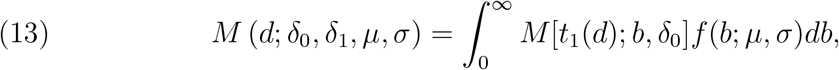

Thus, the formula for the metastasis detection probability depends on the set of model parameters {*δ*_0_, *δ*_1_, *μ*, *σ*}. Metastasis detection does not depend on parameter a, since we assume the screening for metastases occurs at diagnosis, before the treatment that could remove metastases.

### Quantile time to death from cancer for patients with given tumor size

To model any quantile (e.g., median) time it takes to die of cancer, we again make the assumption, that cancer death is due to metastases. We consider the q-th quantile time to death of cancer for patients who (a) had surgery following the diagnosis with diameter *d*, and (*b*) will indeed die of cancer. To this end, we need to derive the time *x*_*q*_ ≤ *t*(*d*) that elapses from tumor onset at which the *q*-th fraction of patients diagnosed with tumor diameter *d* develop metastases that were later not removed by post-surgical treatment.

We first consider the conditional cumulative distribution function of *x*_*q*_, for the waiting time *τ* for metastases, conditioned on that (i) up to time *t*_0_(*d*) = *t*(*d*) − *δ*_0_ no metastasis has been created, and that (ii) the metastases were present at *t*(*d*). For patients with a given value of the bottleneck severity *b*, the form of such conditional distribution function is 
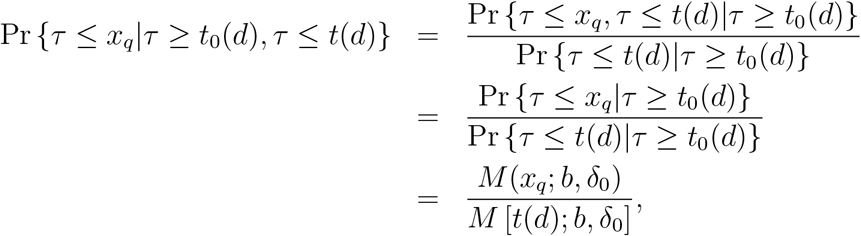
where the last line follows from Equation 4. To account for the variability of the bottleneck, as above we use the marginal probability with respect to the log normal distribution of *b* with parameters *μ* and *σ*. In addition, we use the post-treatment metastasis probabilities, assuming, again as above, that treatment may remove some of the life-threatening metastases. We thus find *x*_*q*_ as the root of the equation 
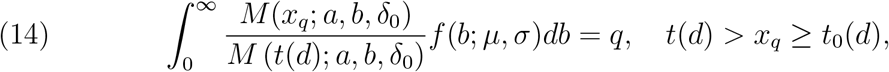

The quantile time to death is then given by 
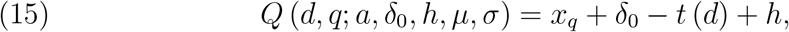
where *h* is an additional treatment-related parameter that accounts for the increase of patient survival due to therapy after detection of the tumor. Thus, the formula for the quantile time to death depends on the cancer-specific parameters {*a*, *δ*_0_, *h*, *μ*, *σ*}, where the dependence on a, *μ*, and *σ* is introduced via *x*_*q*_.

### Quantile time to death from cancer for the subset of patients with detected metastases at diagnosis with given tumor size

For the subset of patients with detected metastases, the quantile time to death should be com-puted conditioning on the fact that the detectable metastases originated before *t*_1_(*d*) = *t*(*d*) − *δ*_1_ and after *t*_0_(*d*) = *t*(*d*) − *δ*_0_. Reasoning analogously to above, we solve 
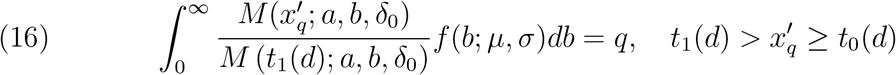
for *x*′_*q*_, and the quantile time to death for patients with metastases detectable at diagnosis is then given by 
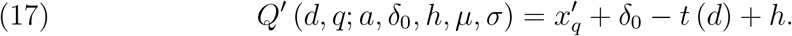

### Model fitting

The model was fitted to metastasis detection rate and quantile times to death for different tumor diameters, obtained from the SEER database, using nonlinear least squares with the Levenberg-Marquardt algorithm implemented in the minpack.lm R package. The optimization was initialized with a total of 2888 different starting values for the parameters, which were sampled from intervals of sensible values: *δ*_0_ ∈ [0,inf], *δ*_1_ ∈ [0,inf], *h* ∈ [0,inf], *μ* ∈ [0.1,inf], *σ* ∈ [0.001,inf], *σ* ∈ [0,1.09], and satisfied the condition *δ*_1_ ≤ *δ*_0_. The upper bound of 1.09 for parameter a corresponds to an upper bound for the probability of removing metas-tases by treatment (Equation 9). Here, we assume that metastasis which originated 5 years before the diagnosis can be removed with probability at most 1%.

The quantile times to death were computed from the data for eleven different quan-tiles, from the 0.35-th to 0.65-th quantile. Using quantile time to death data for multiple quantiles (instead only one quantile, e.g. just median) brought power for model fit from a large dataset.

### Validation on independent dataset

For each of the two pancreatic cancer cohorts, Autopsy and Adjuvant, published by Haeno *et al*. (35), we selected patients who had surgery and died of cancer. Next, for each cohort separately, using their clinical data, we computed the survival curves and collected the diameters of patients. For these diameters, we predicted the quantile times to death for quantiles from 0.05 to 0.95 by 0.05, from our model for pancreatic cancer fitted to SEER data, using Equation 15. Finally, to each quantile we mapped the mean predicted time to death for that quantile across the diameters in the cohort. The prediction of the survival function was obtained as one minus the inverse of this mapping. The analysis performed in (35) did not provide the survival functions for the cohorts directly. To obtain the survival function for the model that the authors (35) fit to the Autopsy cohort, we computed cumulative sum of the frequencies predicted by that model for consecutive survival times, read off from Figures 2A,B in (35). Similarly, the survival function that the same model predicts for the Adjuvant cohort was obtained as cumulative sum of the frequencies plotted in Figures 3B,C,D in (35).

### Predicting impact of treatment decisions

To estimate the marginal impact of surgery delay *δ* for a given tumor size *d*, we compute the difference of cancer death probabilities (Equation 11) with and without the delay 
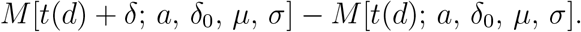

Similarly, for a fixed bottleneck severity *b*, the impact of surgery delay is evaluated using Equation (10) as 
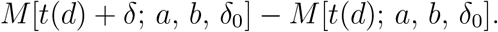

Next, we model increase of chemotherapy efficiency as an increase of the average metastasis age when the metastasis becomes irremovable by treatment, i.e., by multiplying the parameter a by a factor *c*_1_ > 1. To quantify the marginal impact of increased chemotherapy efficiency, we compare the cancer death probabilities with and without the increase, 
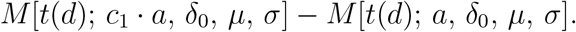

Similarly, for a fixed bottleneck severity *b*, the impact of increased chemotherapy efficiency is evaluated as 
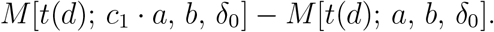

Finally, we predict the impact of increase of the median of the bottleneck distribution by a factor *c*_2_ > 1. Such an increase would correspond to the mechanism of action of vaccines, where we would be able to strengthen the defense of the immune system against the initiation of the metastatic colonies. Since the median of the log-normal distribution is given by exp(*μ*), the new distribution of the bottleneck parameter has location parameter log(*c*_2_) + *μ*. The marginal impact of increase of bottleneck median is calculated comparing cancer death probability with and without the increase 
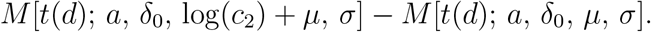

For a fixed bottleneck severity *b*, the impact is evaluated as the difference between the cancer death probability for increased b and for *b* unchanged 
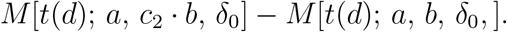

Figure 5 features predictions for *δ* = 16 weeks (0.3 years), *c*_1_ = 1.2, and *c*_2_ = 1.2. In Figure S3 we compare the results for *δ* = 16 weeks with *δ* = 8 weeks, as well as for *c*_1_ = 1.2 with c_1_ = 1.1, and for *c*_2_ = 1.2 with *c*_2_ = 1.1.

### Code availability

The code allowing full reproducibility of all presented results is freely available at https://github.com/EwaSzczurek/MetastaticBottleneck.

## 1. Supplementary Tables

**Table S1.** Overview of analyzed cancer types with minimum follow up times and number of cases included after filtering from the SEER (37) database. For each cancer type, we provide a short name that is used throughout the manuscript. For each type, we defined a designated minimum follow-up time that ensures a sufficient number of patients for calculating representative statistics of frequency of metastasis detection, frequency of cancer death and quantile times to death. Generally, the minimum follow up times were a compromise between the requirement of long monitoring time after diagnosis, and the resulting sample sizes for each cancer. The minimum follow-up time differs between different cancer types, reflecting their clinical behavior. For example, for more aggressive cancers, such as pancreatic and esophageal cancer, which have a poor prognosis, a shorter minimum follow-up time was allowed. In contrast, for breast cancer, where the ten year survival rate for all stages is nearly 80%, the minimum follow-up was set to 20 years. This still allowed for a very large sample size of 35949 patients, since breast cancer is one of the most common cancer types.

| Cancer type<br>short name | Cancer type | Min. follow up<br>in years | Number of cases<br>included |
| --- | --- | --- | --- |
| Breast | Invasive ductal carcinoma | 20 | 35949 |
| Breast lob | Invasive lobular carcinoma | 15 | 6532 |
| Ovarian | Epithelial ovarian cancer | 15 | 2370 |
| Endometrial | Endometrial carcinoma | 15 | 6941 |
| Esophageal | Esophageal carcinoma | 10 | 1725 |
| Gastric | Gastric adenocarcinoma | 10 | 8238 |
| Colon | Adenocarcinoma of the colon<br>(except mucinous) | 15 | 45538 |
| Colon muc | Mucinous adenocarcinoma<br>of the colon | 15 | 3739 |
| Rectal | Adenocarcinoma of the rectum | 15 | 8334 |
| Pancreatic | Exocrine pancreatic cancer | 10 | 2618 |
| Lung | Non-Small cell lung cancer | 15 | 18587 |
| Head & neck | Head and neck<br>squamous-cell carcinoma (HNSCC) | 15 | 8046 |
| Renal | Renal cell carcinoma | 15 | 8133 |
| Bladder | Urothelial carcinoma | 15 | 2441 |

**Table S2.** Significance analysis of the monotonic dependence of clinical variables on tumor size. For each cancer type (rows), the p-values in a two-sided Mann-Kendall trend test are reported. Table entries in red contain p-values lower than 0.01. Light red entries contain p-values lower than 0.05. For metastasis detection probability, cancer death probability and median time to death, the monotonic trend is significant for all cancers, except ovarian and with exception of cancer death probability for esophageal and median time to death for colon mucinous cancer. The same results hold for false discovery rate (FDR) computed from p-values using Benjamini-Hochberg correction for 14 tests (first four columns) or without correction (last four columns). NA values in the “Median time to death for patients with mets” column indicate that, due to too small patient sample sizes, there is no such data available.

| Cancer | FDR (Benjamini-Hochberg adjusted p-values) |  |  |  | p-values |  |  |  |
| --- | --- | --- | --- | --- | --- | --- | --- | --- |
|  | Met detection prob. | Cancer death prob. | Median time to death | Median time to death with mets | Met detection prob. | Cancer death prob. | Median time to death | Median time to death with mets |
| Breast | 8.19e-07 | 3.84e-06 | 3.21e-06 | 1.54e-01 | 3.28e-08 | 6.91e-07 | 3.85e-07 | 1.33e-01 |
| Breast lob | 1.15e-05 | 6.74e-05 | 5.40e-04 | NA | 3.23e-06 | 3.09e-05 | 3.03e-04 | NA |
| Ovarian | 2.56e-01 | 3.81e-02 | 7.07e-01 | 4.86e-01 | 2.30e-01 | 2.97e-02 | 6.78e-01 | 4.48e-01 |
| Endometrial | 7.70e-06 | 7.88e-05 | 6.30e-04 | NA | 1.85e-06 | 3.78e-05 | 3.66e-04 | NA |
| Esophageal | 6.74e-05 | 5.81e-01 | 6.63e-04 | NA | 3.10e-05 | 5.46e-01 | 4.01e-04 | NA |
| Gastric | 7.17e-06 | 1.51e-05 | 1.51e-05 | 7.34e-01 | 1.58e-06 | 5.12e-06 | 4.89e-06 | 7.19e-01 |
| Colon | 5.82e-06 | 3.84e-06 | 6.04e-07 | 1.49e-02 | 1.16e-06 | 6.28e-07 | 1.21e-08 | 1.10e-02 |
| Colon muc | 5.08e-04 | 4.24e-02 | 7.85e-01 | 2.02e-01 | 2.74e-04 | 3.39e-02 | 7.85e-01 | 1.78e-01 |
| Rectum | 3.84e-06 | 1.44e-06 | 1.40e-05 | 1.36e-01 | 6.28e-07 | 8.64e-08 | 4.19e-06 | 1.14e-01 |
| Pancreas | 1.60e-05 | 2.57e-02 | 1.19e-04 | NA | 5.76e-06 | 1.95e-02 | 5.96e-05 | NA |
| Lung | 5.00e-03 | 6.63e-04 | 3.21e-06 | 1.03e-02 | 3.40e-03 | 4.11e-04 | 3.19e-07 | 7.43e-03 |
| Head & neck | 1.75e-04 | 3.15e-05 | 8.46e-06 | NA | 9.10e-05 | 1.26e-05 | 2.20e-06 | NA |
| Renal | 3.21e-06 | 6.40e-05 | 5.63e-03 | 8.72e-02 | 3.26e-07 | 2.69e-05 | 3.94e-03 | 7.15e-02 |
| Bladder | 2.68e-05 | 1.65e-03 | 9.20e-04 | NA | 1.02e-05 | 1.09e-03 | 5.89e-04 | NA |

**Table S3.** Per-cancer tumor growth rates fixed for thirteen modeled cancer types. For each cancer type, we list the tumor doubling time in days, denoted *D*, as found reported (usually as median or mean value across a measured population) in the cited references. The per year exponential growth rate r in our model, was then computed from those doubling times as *D* · ln(2)/365.24. For several cancer types, we were unable to find measured doubling times. For those cancer types, we fixed the corresponding rates to ones measured in the most similar cancer type. Specifically, for endometrial cancer, we fixed the doubling rate to the rate reported in another female cancer, breast. For esophageal, gastric and bladder cancer types, we fixed the rate as reported for colon and rectum.

| Cancer | Doubling time $D$ (days) | Reference |
| --- | --- | --- |
| Breast | 212 | (42) |
| Breast lob | 212 | (42) |
| Endometrial | 212 | (42) |
| Esophageal | 255 | (43) |
| Gastric | 255 | (43) |
| Colon | 255 | (43) |
| Colon muc | 255 | (43) |
| Rectal | 255 | (43) |
| Pancreatic | 144 | (44) |
| Lung | 166,3 | (45) |
| Head & neck | 99 | (46) |
| Renal | 603 | (47) |
| Bladder | 255 | (43) |

## 2. Supplementary Figures

**Figure S1.**
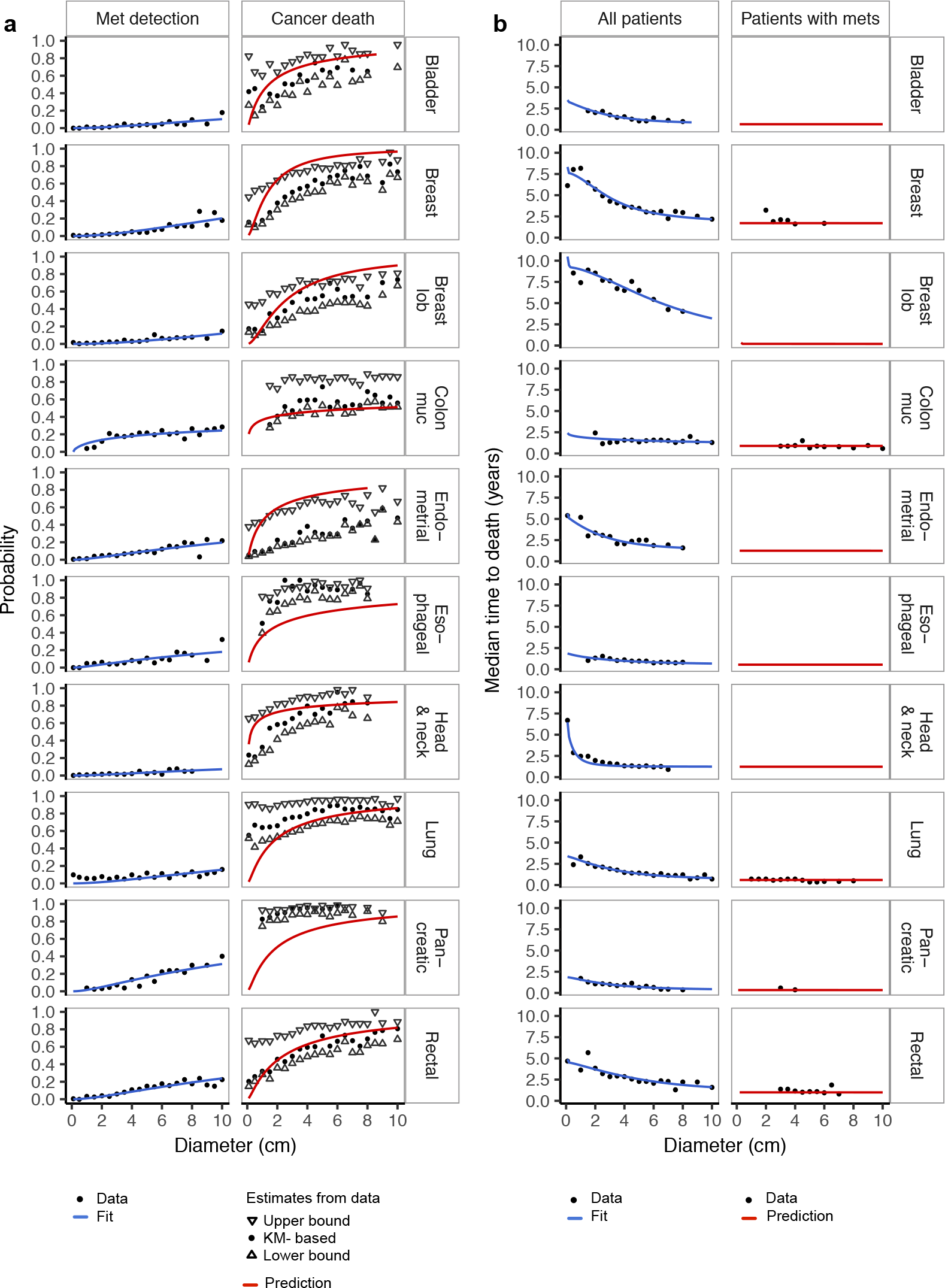
Fit and validation of the model on the SEER data, for ten cancer types (rows, alphabetically ordered), not featured in Figure 3. Black points represent data records, blue lines show fitted, and red lines predicted curves. The model fits the data perfectly. **a** For cancer death probability, up and down-oriented triangles present the upper and lower estimates of that variable from the data, respectively, while the black dots represent a Kaplan-Meier (KM) derived estimate (Methods). In agreement with the data, we model cancer death frequency (right panel) as the probability of metastasis which were not removed by treatment, which accounts also for undetectable metastases and is thus larger than metastasis detection probability (left panel). **b** Despite the fact that the median time to death data for all patients (left panel), which was used for model fitting is tumor-size dependent, the model correctly predicts that median time to death for patients with metastases detected at diagnosis (right panel) is constant across tumor diameters.

**Figure S2.**
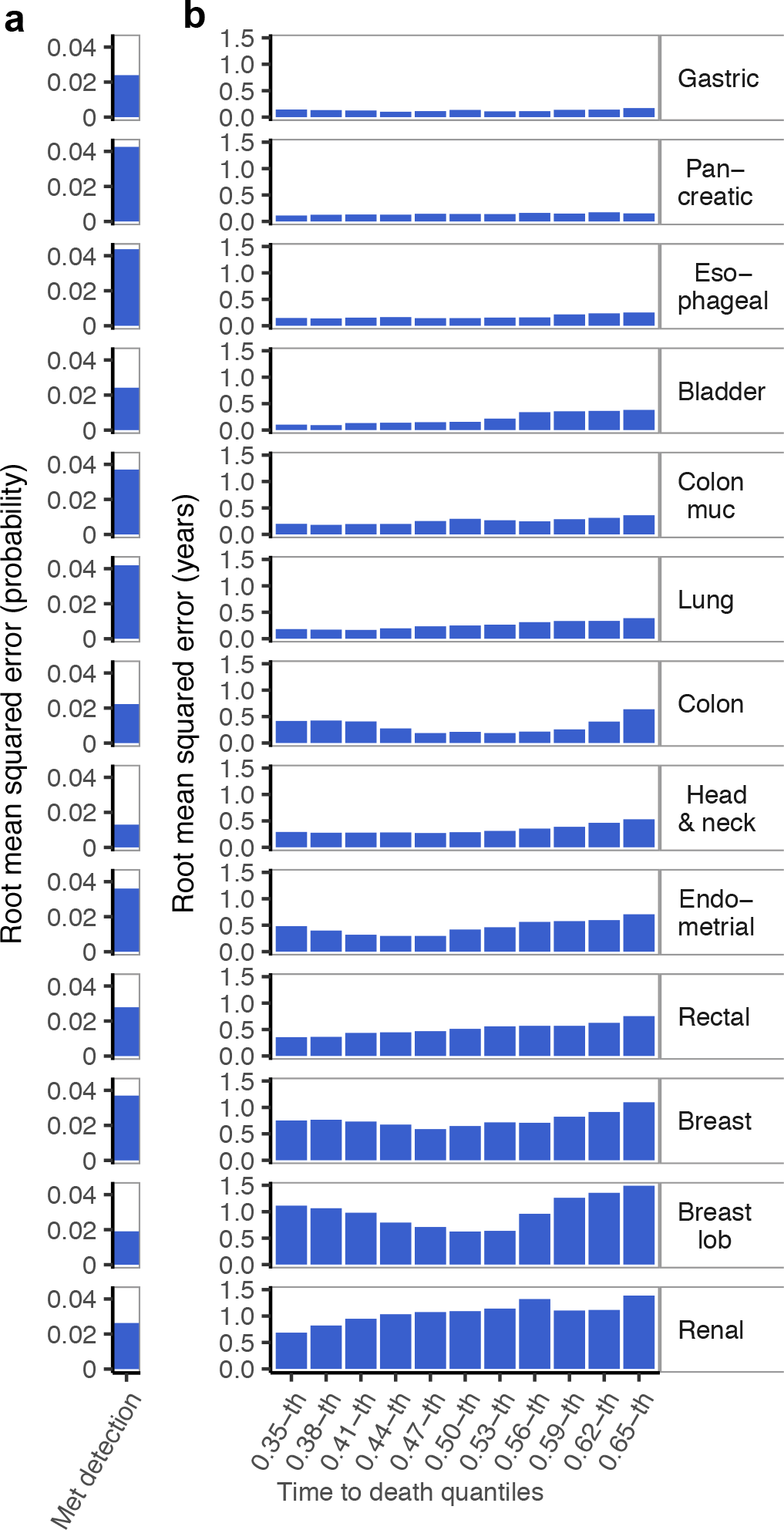
Summary of the model fit. For thirteen different cancer types, their fit to the data is measured using root mean squared error (RMSE). The cancer types (rows) are ordered by the increasing mean RMSE of the fit to quantile time to death data. **a** Low RMSE values indicate very good agreement of the model with metastasis detection probability (as visualized also in Figure 3a and Figure S1a, blue lines). The largest RMSE is obtained for pancreatic cancer, for which also the largest absolute metastasis detection probability is recorded (see Figure S1a). **b** Similarly good fit is obtained for quantile time to death data, for eleven different quantiles (x-axis). Larger RMSE values than for metastasis detection probability in (**a**) are due to the fact that quantile time to death is in measured in (usually, several) years. The fit to 0.5-th quantile (median) is visualized also in Fig. 3b and Figure S1b. The RMSE for other quantiles is comparable to the RMSE obtained for the median (0.50-th quantile) time to death data.

**Figure S3.**
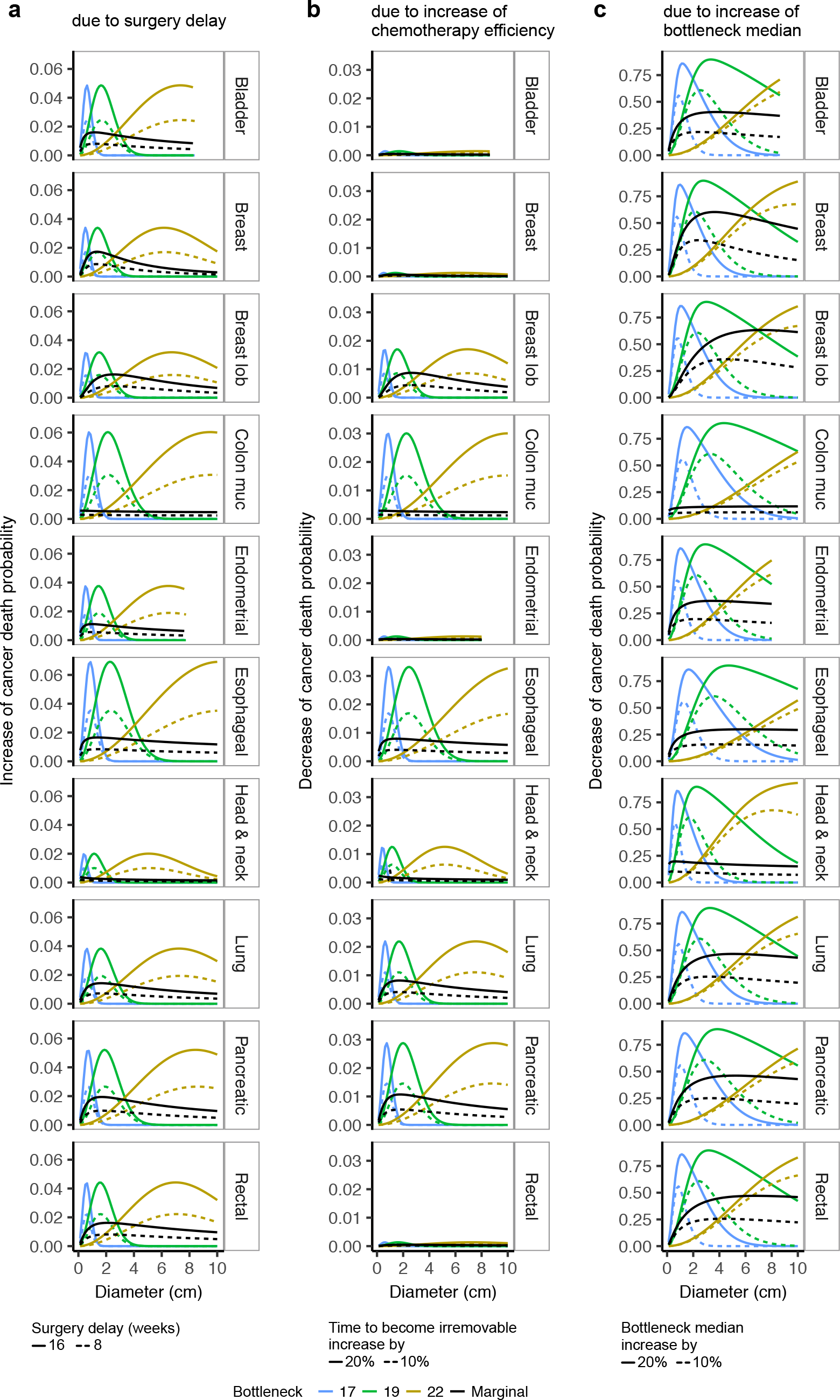
Predicted impact of treatment decisions. Change in cancer death probability (y-axis) due to treatment change depends on tumor diameter (*x*-axis) and bottleneck severity (colors), shown for ten cancers not featured in Figure 4 in the main text (rows; ordered alphabetically). Black curves present the change in cancer death probability marginalized over the bottleneck severity. The increase of cancer death probability due to surgery delay by either 16 or 8 weeks (solid or dashed lines in **a**, respectively), as well as decrease of cancer death probability due to increase of chemotherapy efficacy by 20% (solid lines in **b**) or by 10% (dashed lines in **b**), are both much smaller than the decrease of cancer death probability due to increase of the metastatic bottleneck median by 20% (solid lines in **c**) or by 10% (dashed lines in **c**).

### Derivation of the metastasis success probability

Consider the metastasis initiation model introduced in the main text, where for colony size *i* the proliferation rate is *λ*_*i*_ and death rate is *μ*_*i*_ 
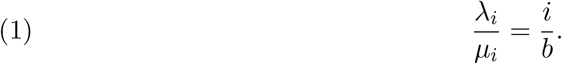

We will now show that for this model the metastasis success probability *s*_*i*_, equal to the probability that the colony starting from single cell will not go extinct, is given by 
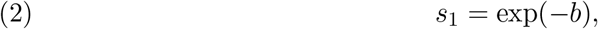
while the probability of colony survival starting from *i* > 1 cells equals 
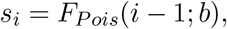
where *F*_*Pois*_(*i*; *b*) denotes a cumulative Poisson distribution function with parameter *b*, evaluated at value *i*.

In a birth and death chain with a single absorbing state *i* = 0, the probability of success of the metastasis starting from state *i* can be computed with the recursion (1,2) 
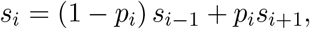
where *p*_*i*_ denotes the probability of jumping from the state of *i* to *i* + 1 cells 
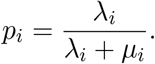

Setting 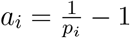, with *s*_0_ = 0, and rearranging terms we obtain 
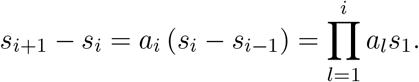

Hence, 
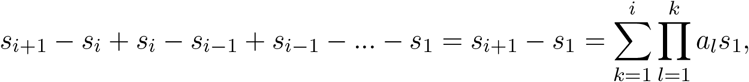

yielding 
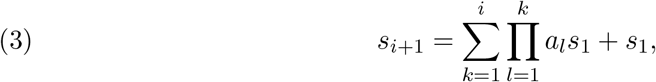
and, since *s*_*i*_ → 1 as *i* → ∞, 
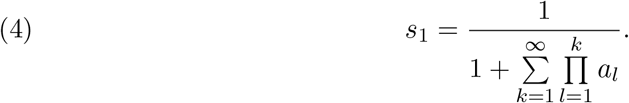

With the birth rate satisfying 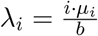 we have 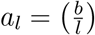 and therefore 
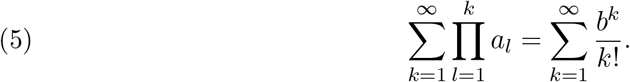

This together with Equation (4) gives 
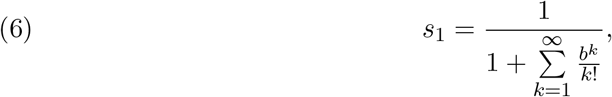
and together with Equation (5) yields the probability of survival from a given state *i* > 1 
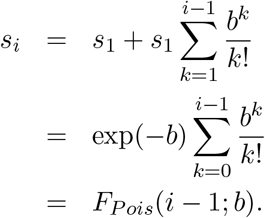

Finally, from the Taylor series 
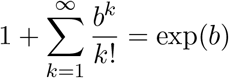
and Equation (6) we obtain the metastasis success probability 
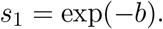

### Discussion of timing of the stochastic process of metastasis initiation

The assumption of the birth to death ratio satisfying Equation (1) does not define the timing of events in the process. The assumption is enough to derive the probabilities of metastatic success, which, as shown above, do not depend on the particular values of birth λ_*i*_ and death rates *μ*_*i*_.

To describe the timing of this process, a convenient way would be to set 
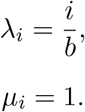

In such a way, we could model the real time behavior of the process, where the time unit would be the mean time to death. Thus, realistic time predictions would depend on the measurement of average time to death of cells in a growing metastatic colony.

### Consideration of truncated birth to death rates ratio

We consider a different assumption, namely that 
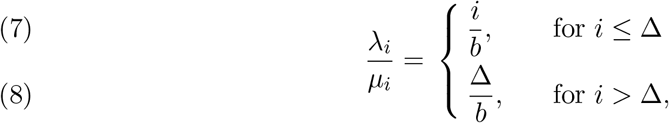
where Δ is a threshold value satisfying Δ > *b*, and let 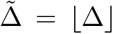. We will refer to the stochastic metastasis initiation model with this assumption as the truncated model. The survival probability starting with a single cell in the truncated model is denoted by *s*_1,Δ_ Clearly, we have *s*_1,Δ_ < *s*_1_, where *s*_1_ is the survival probability derived above for the stochastic model without the truncation.

With this assumption, Equations (3) and (4) are also satisfied. Moreover, with a defined as above, we obtain 
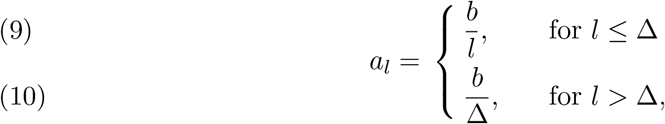
and therefore for the denominator in Equation (4) we get 
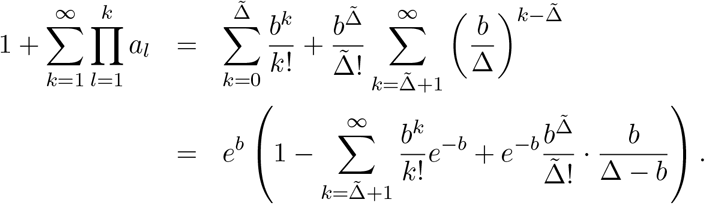

Since *s*_1,Δ_ < *s*_1_, we have 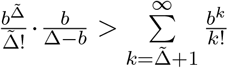 and hence 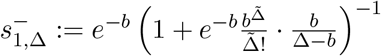 is a lower bound to *s*_1,Δ_. Defining 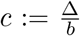 and 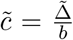 we obtain with the lower bound for the factorial 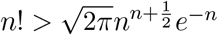 
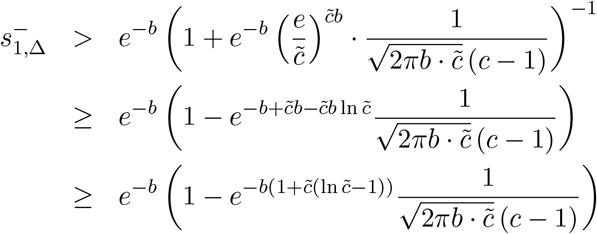

Note that 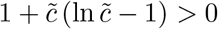 for 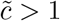. For 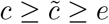 we obtain a simple uniform lower bound, namely 
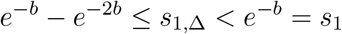
but already for smaller values of c the difference between s_1_ and *s*_1;Δ_ can essentially be neglected. As an example let *b* = 10 and 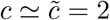. We obtain 
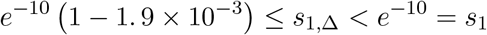

Hence the error made by replacing the survival probability for the truncated process by the one of the process without truncation is less then two thousand parts in this example.

### Generalization of the metastasis initiation model

In the general case, we will consider the following assumption about the ratio of rates λ_*i*_ and *μ*_*i*_ for cell division and death, respectively: 
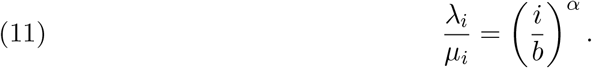

Here, the exponent *α* defines the nature of the dependence of proliferation to death rate ratio on colony size. We set *α* = 1, as in the main text, to define linear dependence. Another choice of *α* = 1/3 would correspond to the assumption that the proliferation rate depends on the ratio of the number of cells on the surface to the total number of cells in the colony, assuming spherical metastasis growth (see below). For a population of cells of size *i*, the individual rates λ_*i*_ and *μ*_*i*_ correspond to population proliferation rate *i* · λ_*i*_ = *i*^*a*+1^/(*b*)^*α*^ · *μ*_*i*_ and population death rate *i* · *μ*_*i*_, respectively. We will now derive approximation for the metastasis survival probability in this general case.

Let *α* = 1. Following the same argument as above, we can derive 
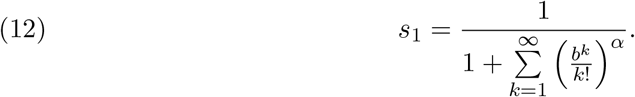

We have: 
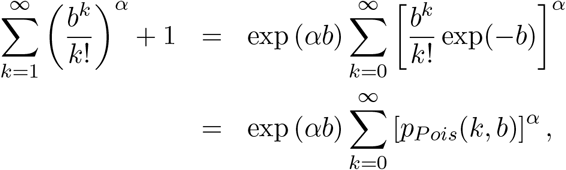
where 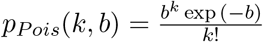 is the Poisson distribution with parameter *b*.

Jacquet *et al*. (3) showed that for such sequences of distributions *p*(*k*, *n*), which can be approximated with a normal distribution with variance *nσ*^2^ > 0 for *n* → ∞, the Renyi entropy satisfies 
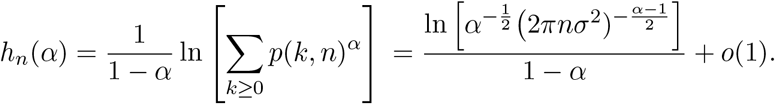

The Poisson distribution with parameter *b* sufficiently large is well approximated by the the normal distribution with variance *b* (and mean *b*). Thus, for the Poisson distribution with rate *b* its Renyi entropy, denoted (*b*), becomes 
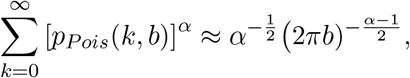

Simplifying, we obtain 
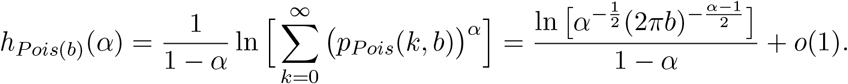
which leads to 
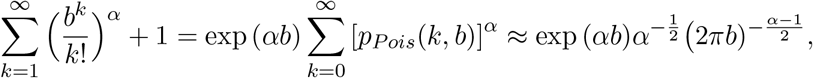
and after inserting into Equation 12 the metastasis success probability in this model becomes 
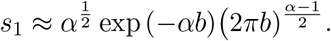

Note that for *α* = 1, in agreement with Equation 2, the right hand side equals exp(-*b*).

To consider a possible biologically relevant value of *α* ≠ 1, we discuss a scenario where the forming colony is a sphere and where, out of all cells in the colony, the cells on the sphere surface are most vulnerable, due to exposure to the surrounding environment.

We can further approximate that the total number of cells i is on the order of the volume of the sphere, *i* = *O*(*ρ*^3^), and the number of the cells on its surface is *j* = *O*(*ρ*^2^), where *ρ* denotes the radius of the sphere. Thus, *j* = *O*(*i*^2/3^). Assuming that the proliferation rate of cells in the colony depends on the ratio *i*/*j* = *i*^1/3^ between the total to the number of cells on the surface we obtain 
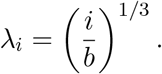

Thus, for spherical metastases, the model parameter is 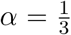 and the survival probability becomes 
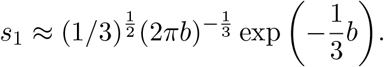

## References

[1] Fidler, I. J. The pathogenesis of cancer metastasis: the ‘seed and soil’ hypothesis revisited. Nature Rev. Cancer 3, 453–458 (2003).

[2] Sporn, M. B. The war on cancer. Lancet 347, 1377–1381 (1996).

[3] Hanahan, D. & Weinberg, R. A. Hallmarks of cancer: the next generation. Cell 144, 646–674 (2011).

[4] Fidler, I. Metastasis: guantitative analysis of distribution and fate of tumor embo-lilabeled with 125 i-5-iodo-2’-deoxyuridine. Journal of the National Cancer Institute 45, 773–82 (1970).

[5] Weiss, L. Metastatic inefficiency. Adv. Cancer Res. 54, 159–211 (1990).

[6] Chambers, A. F., Groom, A. C. & MacDonald, I. C. Dissemination and growth of cancer cells in metastatic sites. Nature Rev. Cancer 2, 563–572 (2002).

[7] Chaffer, C. L. & Weinberg, R. A. A perspective on cancer cell metastasis. Science 331, 1559–1564 (2011).

[8] Nguyen, D. X., Bos, P. D. & Massague, J. Metastasis: from dissemination to organ-specific colonization. Nature Rev. Cancer 9, 274–284 (2009).

[9] Nguyen, D. X. & Massague, J. Genetic determinants of cancer metastasis. Nature Rev. Genet. 8, 341–352 (2007).

[10] Liu, W. Copy number analysis indicates monoclonal origin of lethal metastatic prostate cancer. Nature Med. 15, 559–565 (2009).

[11] Navin, N. Tumour evolution inferred by single-cell sequencing. Nature 472, 90–94 (2011). URL http://dx.doi.org/10.1038/nature09807.

[12] Hosseini, H. et al. Early dissemination seeds metastasis in breast cancer. Nature (2016).

[13] Harper, K. L. et al. Mechanism of early dissemination and metastasis in Her2(+) mammary cancer. Nature (2016).

[14] Poste, G. & Fidler, I. J. The pathogenesis of cancer metastasis. Nature 283, 139–146 (1980).

[15] Luzzi, K. J. Multistep nature of metastatic inefficiency: dormancy of solitary cells after successful extravasation and limited survival of early micrometastases. Am. J. Pathol. 153, 865–873 (1998).

[16] Cameron, M. D. Temporal progression of metastasis in lung: cell survival, dormancy, and location dependence of metastatic inefficiency. Cancer Res. 60, 2541–2546 (2000).

[17] Varghese, H. J. Activated ras regulates the proliferation/apoptosis balance and early survival of developing micrometastases. Cancer Res. 62, 887–891 (2002).

[18] Wong, C. W. et al. Apoptosis: An early event in metastatic inefficiency. Cancer Research 61, 333–338 (2001).

[19] Paget, S. The distribution of secondary growths in cancer of the breast. 1889. Cancer Metastasis Rev. 8, 98–101 (1989).

[20] Langley, R. R. & Fidler, I. J. The seed and soil hypothesis revisited-the role of tumor-stroma interactions in metastasis to different organs. Int. J. Cancer 128, 2527–2535 (2011).

[21] Liotta, L. A., Saidel, M. G. & Kleinerman, J. The significance of hematogenous tumor cell clumps in the metastatic process. Cancer Res. 36, 889–894 (1976).

[22] Aceto, N. et al. Circulating tumor cell clusters are oligoclonal precursors of breast cancer metastasis. Cell 158, 1110–1122 (2014).

[23] Brierley, J. D., Gospodarowicz, M. K. & Wittekind, C. e. (eds.) TNM Classification of malignant tumours (2017), 8th edn.

[24] Koscielny, S. Breast cancer: relationship between the size of the primary tumour and the probability of metastatic dissemination. Br. J. Cancer 49, 709–715 (1984).

[25] Engel, J. et al. The process of metastasisation for breast cancer. Eur. J. Cancer 39, 1794–1806 (2003).

[26] Liotta, L. A., Kleinerman, J. & Saidel, G. M. Quantitative relationships of in-travascular tumor cells, tumor vessels, and pulmonary metastases following tumor implantation. Cancer Res. 34, 997–1004 (1974).

[27] Saidel, G. M., Liotta, L. A. & Kleinerman, J. System dynamics of metastatic process from an implanted tumor. J. Theor. Biol. 56, 417–434 (1976).

[28] Liotta, L. A., Saidel, G. M. & Kleinerman, J. Stochastic model of metastases formation. Biometrics 32, 535–550 (1976).

[29] Iwata, K., Kawasaki, K. & Shigesada, N. A dynamical model for the growth and size distribution of multiple metastatic tumors. J. Theor. Biol. 203, 177–186 (2000).

[30] Hanin, L., Rose, J. & Zaider, M. A stochastic model for the sizes of detectable metastases. J. Theor. Biol. 243, 407–417 (2006).

[31] Hartung, N. et al. Mathematical modeling of tumor growth and metastatic spreading: validation in tumor-bearing mice. Cancer Res. 74, 6397–6407 (2014).

[32] Michor, F., Nowak, M. A. & Iwasa, Y. Stochastic dynamics of metastasis formation. J Theor Biol 240, 521–530 (2006).

[33] Dingli, D., Michor, F., Antal, T. & Pacheco, J. M. The emergence of tumor metastases. Cancer Biol. Ther. 6, 383–390 (2007).

[34] Haeno, H. & Michor, F. The evolution of tumor metastases during clonal expansion. J Theor Biol 263, 30–44 (2010).

[35] Haeno, H. et al. Computational modeling of pancreatic cancer reveals kinetics of metastasis suggesting optimum treatment strategies. Cell 148, 362 – 375 (2012).

[36] Benzekry, S. et al. Modeling Spontaneous Metastasis following Surgery: An In Vivo-In Silico Approach. Cancer Res. 76, 535–547 (2016).

[37] SEER. Surveillance, Epidemiology, and End Results Program (www.seer.cancer.gov) Research Data (1973–2013), National Cancer Institute, DCCPS, Surveillance Research Program, Surveillance Systems Branch (released in Nov 2015).

[38] Lengyel, E. Ovarian cancer development and metastasis. Am. J. Pathol. 177, 10531064 (2010).

[39] DeVita, V. T., Young, R. C. & Canellos, G. P. Combination versus single agent chemotherapy: a review of the basis for selection of drug treatment of cancer. Cancer 35, 98–110 (1975).

[40] Melero, I. et al. Therapeutic vaccines for cancer: an overview of clinical trials. Nat Rev Clin Oncol 11, 509–524 (2014).

[41] Gerlee, P. The model muddle: in search of tumour growth laws. Cancer Research (2013).

[42] von Fournier, D. et al. Growth rate of 147 mammary carcinomas. Cancer 45, 21982207 (1980).

[43] Choi, S. J., Kim, H. S., Ahn, S. J., Jeong, Y. M. & Choi, H. Y. Evaluation of the growth pattern of carcinoma of colon and rectum by MDCT. Acta Radiol 54, 487–492 (2013).

[44] Furukawa, H., Iwata, R. & Moriyama, N. Growth rate of pancreatic adenocarcinoma: initial clinical experience. Pancreas 22, 366–369 (2001).

[45] Arai, T. et al. Tumor doubling time and prognosis in lung cancer patients: evaluation from chest films and clinical follow-up study. Japanese Lung Cancer Screening Research Group. Jpn. J. Clin. Oncol. 24, 199–204 (1994).

[46] Jensen, A. R., Nellemann, H. M. & Overgaard, J. Tumor progression in waiting time for radiotherapy in head and neck cancer. Radiother Oncol 84, 5–10 (2007).

[47] Ozono, S. et al. Tumor doubling time of renal cell carcinoma measured by CT: collaboration of Japanese Society of Renal Cancer. Jpn. J. Clin. Oncol. 34, 82–85 (2004).

## REFERENCES

[1] K.B. Athreya and P.E. Ney. Branching processes. Dover, Mineola, New York, 1972.

[2] P. Haccou, P. Jagers, and V. A. Vatutin, editors. Branching processes: Variation, growth, and extinction of populations. Cambridge University Press, 2005.

[3] Jacquet Philippe, Szpankowski Wojciech, and Ln N. Entropy computations via analytic depoissonization. IEEE Trans. Information Theory, 45:1072–1081, 1998.

